# Role of Immature Choroid Plexus in the Pathology of Autism Spectrum Disorder

**DOI:** 10.1101/2023.12.01.569662

**Authors:** Motoi Tanabe, Yuga Saito, Ayaka Takasaki, Keita Nakano, Chikako Suzuki, Nao Kawamura, Aki Hattori, Shun Nagashima, Mami Oikawa, Shigeru Yanagi, Tomoyuki Yamaguchi, Toshifumi Fukuda

## Abstract

During gestation, the choroid plexus (ChP) produces protein-rich cerebrospinal fluid and matures prior to brain development. It is assumed that ChP dysfunction has a profound effect on developmental neuropsychiatric disorders, such as Autism Spectrum Disorder (ASD). However, the mechanisms linking immature ChP to the onset of ASD remain unclear. In this study, we found that ChP-specific CAMDI-knockout mice developed an immature ChP, alongside decreased multiciliogenesis and expression of differentiation marker genes following disruption of the cerebrospinal fluid barrier. These mice exhibited ASD-like behaviors, including impaired socialization with delayed critical period. Additionally, administration of Metformin, an FDA-approved drug, before the critical period achieved ChP maturation and restored social behaviors. Furthermore, ASD model mice and ASD patient-derived organoids developed immature ChP. These results indicate towards involvement of immature ChP in the pathogenesis of ASD and suggest the targeting of functional maturation of ChP as a therapeutic strategy for ASD.

## INTRODUCTION

The incidence of neurodevelopmental and neuropsychiatric disorders such as Autism Spectrum Disorder (ASD) has seen a rise among children and adolescents in recent years. Often these disorders are associated with poor quality of life due to congenital brain dysfunction.^1,2^ These neurodevelopmental disorders are usually functional or organic and several genetic factors are thought to play a role in their pathogenesis. Factors such as neuronal proliferation, differentiation, and migration; synapse formation; imbalance between excitatory and inhibitory neurons; monoaminergic neuronal differentiation; and increased immune mechanisms in the brain including microglial and astrocyte activation, are thought to contribute to the pathogenesis of neurodevelopmental and neuropsychiatric disorders.^3–7^ In addition, dysfunction of normal plasticity during the critical period has been identified as a contributing factor in neurodevelopmental and neuropsychiatric disorders.^8–10^ ASD is characterized by severe deficits of socialization, communication, and repetitive behaviors,^11^ suggesting that alteration of the critical period contribute to these behaviors.^12^ However, the development of fundamental treatments based on pathogenic mechanisms of disorders such as ASD has not yet been achieved, bringing an urgency to the research and development of such treatments.

The choroid plexus (ChP) is a vascular-rich tissue composed of multiciliated epithelial cells differentiated from ependymal cells on ventricular surfaces.^13^ It is the main tissue responsible for the production of cerebrospinal fluid (CSF), and contributes to the interference of water content and shape retention in the brain.^14^ Transthyretin (TTR) is a differentiation marker gene of the ChP and acts as a carrier for the thyroid hormone thyroxine (T4) and retinol.^15–18^ Reduced TTR expression in patients with schizophrenia and depression has been reported.^19–21^ In addition, visual experience promotes the accumulation of non-cell-autonomous Otx2 in parvalbumin (PV)-expressing cells and accelerates the timing of the critical period.^22^ Furthermore, it has been suggested that abnormalities in the critical period may lead to ASD symptoms and other developmental disorders.^12,23^ It was reported that the CSF in the week before and after birth contains high concentrations of proteins.^24^ These reports suggest that changes in the ChP function and CSF composition significantly affect brain development. However, the mechanism linking functionally immature ChP to ASD, particularly to reduced social behavior, remains unclear.

CAMDI (also known as CCDC141) has been identified as a novel binding protein for the schizophrenia-associated protein DISC1,^25^ suggesting CAMDI dysfunction to be closely associated with ASD symptoms.^26–29^ Inhibition of CAMDI expression impairs neuronal migration with unstable centrosome and function in the cerebral cortex during development^25,30–32^ long with GnRH neuronal migration.^33^ Mice lacking the CAMDI gene have shown neuropsychiatric behaviors aligning with symptoms of ASD, such as anxiety, depression, and reduced social behaviors through HDAC6 hyperactivation.^34^ However, it remains unclear whether abnormal neuronal migration can be attributed to development of ASD-like symptoms.

In this study, we showed that impaired multiciliogenesis leads to development of an immature ChP and disruption of the brain-CSF barrier, contributing to reduced social behavior with delay of the critical period. We also demonstrated postnatal administration of the diabetic drug Metformin before the critical period which resulted in ChP maturation and significantly improved social behavior with no delay of the critical period. Furthermore, presence of immature ChP was confirmed in ASD mouse models and ASD patient-derived organoids. These results suggest that a potential novel therapeutic target for ASD is the immature ChP and that treatment before the critical period during postnatal life may be successful.

## RESULTS

### Immature Choroid Plexus and ASD-like behavior in ChP-KO mice

In a previous study, we reported that CAMDI gene-deficient mice exhibit abnormal neuronal migration in the cerebral cortex with immature centrosomes through HDAC6 hyperactivation and showcase ASD-like symptoms such as impaired social behavior.^34^ To elucidate the molecular mechanisms underlying these phenomena, we used DNA microarray technique to comprehensively analyze gene expression in the whole cerebrum of *CAMDI*-deficient mice on postnatal day (P) 0. We found that ChP marker genes, such as TTR, FolR1, Clic6, Cldn2, and Kcne2, were markedly downregulated (Table S1).

We confirmed a high expression signal of CAMDI mRNA in the lateral ventricle ChP on embryonic day (E) 18.5 by in situ hybridization and immunohistochemical (IHC) analysis (Figure 1A). In contrast, FoxJ1 regulates migration of centrioles to the apical surface during formation of multicilia.^35^ It has been reported that FoxJ1 expression is confined to the ChP epithelial cells until E15.5, and that FoxJ1^CreERT2::GFP^ mice have a Cre recombinase inserted at the FoxJ1 locus.^36^ To explore the function of CAMDI in ChP, we generated ChP epithelial cell-specific CAMDI KO (ChP-KO) mice via intraperitoneal administration of tamoxifen on E15.5, obtained by crossing FoxJ1^CreERT2::GFP^ mice with CAMDI flox/flox mice. Quantitative PCR analysis showed that CAMDI mRNA expression was significantly decreased in the ChP of ChP-KO mice (Figure 1B). In contrast, CAMDI mRNA expression and cortical migration remained unchanged in the cerebral cortex (Figures S1A and 1B). Western blot analysis confirmed that CAMDI expression was significantly decreased in the ChP (Figures 1C and 1D). CAMDI regulates centrosome maturation,^34^ which is the basis of ciliogenesis. Fluorescent immunostaining revealed that the ratio of multiciliogenesis was significantly reduced with the abnormal location of replicated centrioles (Figures 1E and 1F). We quantified the expression of FoxJ1, a transcription factor essential for multicilia formation,^37^ and found that IFT88 which is required for protein transport into the cilia^38^ was downregulated (Figures 1G and 1H). Furthermore, the mRNA and protein expression levels of TTR, a differentiation marker in mature ChP epithelial cells, were significantly decreased in ChP-KO mice (Figures 1I–1K). As previously reported, mature ChP epithelial cells express multiciliogenesis genes,^39,40^ our findings revealed downregulation of these genes and TTR, thereby confirming presence of immature ChP epithelial cells in ChP-KO mice.

**Figure 1.**
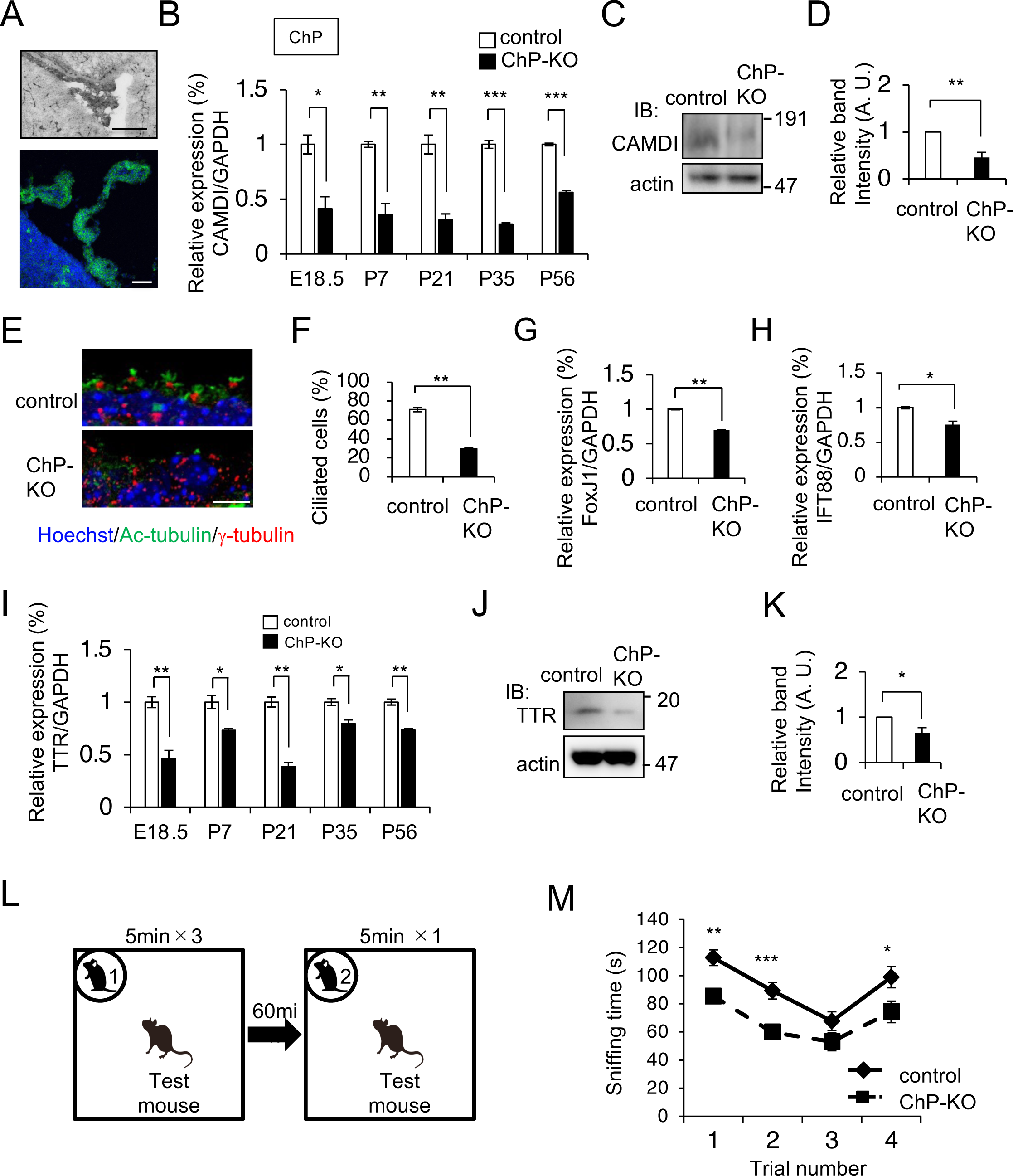
Immature Choroid plexus and ASD-like behavior in ChP-KO mice. A. In situ hybridization and Immunohistochemical analyses of CAMDI in the lateral ventricle ChP at E18.5. Scale bar, 100 μm. B. qRT-PCR analysis of CAMDI mRNA expression in the ChP (normalized to GAPDH). N = 3 mice/group. C. Western blot analysis of CAMDI expression in the ChP at E18.5. D. Quantification of western blots of CAMDI in the ChP (normalized to actin). N = 3 mice/group. E. Immunohistochemical analysis for Ac-tubulin (cilia, green), γ-tubulin (centrosome, red) and Hoechst (nuclear, blue) of the ChP at E18.5. Scale bar, 10 μm. F. Percentage of ciliated (Ac-tubulin-positive) cells of ChP. N = 3 mice/group. G, H. qRT-PCR analysis of FoxJ1 (G) and IFT88 (H) mRNA expression in the ChP at E18.5 (normalized to GAPDH). N = 3 mice/group. I. qRT-PCR analysis of TTR mRNA expression in the ChP at E18.5 (normalized to GAPDH). N = 3 mice/group. J. Western blot analysis of TTR expression in the ChP at E18.5. K. Quantification of western blots for TTR in the ChP (normalized to actin). N = 3 mice/group. L. Schema of Social recognition test. M. Comparison of social recognition time. Time spent sniffing in stimulus mice. Trials 1–3 represent repeated presentations of the same stimulus mouse, and trial 4 represents the introduction of a novel mouse. Control mice (N = 11) and ChP-KO mice (N= 12). *** *P < 0.001*, ** *P < 0.01*, * *P < 0.05*. Data are presented as Mean ± SEM.

Next, we determined whether ChP-KO mice exhibited ASD-like behavior. In the open field test, unlike whole-body KO mice, no differences in hyperactivity, time in the center, or grooming time were observed (Figures S1C–S1E). In the light and dark test, which evaluates anxiety, there was no difference in the latency to enter the light chamber; however, there was a decrease in the time spent in the light chamber, indicating increased anxiety (Figures S1F and S1G). The latency to the first fall from high altitude was significantly shorter, indicating an increase in impulsivity (Figure S1H). ChP-KO mice exhibited repetitive behavior in the marble-burying test, indicating increased compulsive behavior (Figure S1I). In the forced swimming test, ChP-KO mice exhibited increase in immobility time, indicating depression-like behavior (Figure S1J). The hot-plate test revealed hyposensitivity in ChP-KO mice, indicating hypoesthesia (Figure S1K). To investigate whether immature ChP affects social behavior, we performed the three-chamber social interaction test. Although there was no change in time spent in chamber, the sniffing time was significantly reduced in ChP-KO mice (Figures S1L and S1M). In addition, ChP-KO mice were subjected to a social recognition test (Figure 1L). ChP-KO mice showed reduced sniffing times, indicating significantly reduced social behavior (Figure 1M). Altogether, these results indicate that CAMDI contributes to the maturation of ChP epithelial cells, and that its deficiency can cause ASD-like behavior.

### Alteration of multiciliogenesis and inflammation in ChP-KO

To comprehensively analyze the phenomena caused by CAMDI gene deletion in ChP epithelial cells, we verified the changes in gene expression by RNA-seq analysis using the ChP from control and ChP-KO mice at P21. Differentially expressed genes (DEGs) were visualized using a heat-map of four subclusters (Figure 2A). Compared to control mice, ChP-KO mice showed decreased mRNA expression associated with multicilia formation and motility in the ChP (Figure 2B; Table S2). The volcano plot shows DEGs related to multiciliogenesis (Figure 2C). Significant reductions were observed for a group of genes, including FoxJ1, essential for the differentiation of multiciliated cells (Figures 2D and 2E), and Rfx, involved in cilia-related processes (Figure 2F). In addition, there was a significant decrease in the expression of motile cilia-related genes (Figures S2A– 2E).^41^

**Figure 2.**
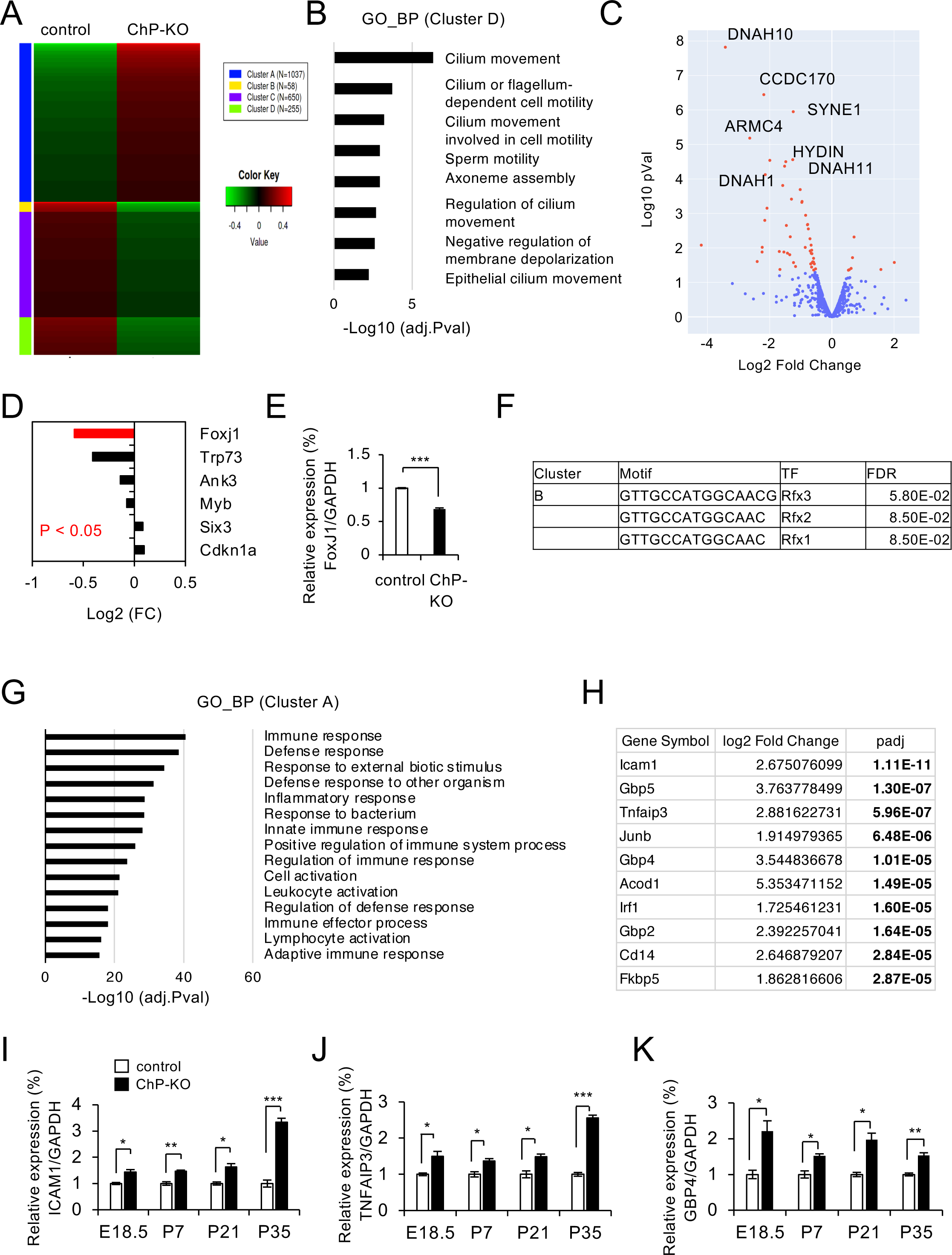
Alteration of multiciliogenesis and inflammation in ChP-KO mice. A. Heatmaps showing differentially expressed transcripts identified by RNA-seq of the choroid plexus in ChP-KO mice at P21. Red indicates high expression while green means low expression of transcripts. B. Enriched GO Biological Process (Cluster D) terms for the downregulated genes at the ChP in ChP-KO mice. C. Volcano plot showing differentially expressed genes at P21 between control and ChP-KO mice. Red: *p* value < 0.05, blue: *p* value > 0.05. D. The expression of the key regulators of multiciliogenesis in ChP of ChP-KO mice. E. qRT-PCR analysis of FoxJ1 expression in ChP-KO mice compared at E18.5. F. Motif enrichment analysis of promoters of downregulated genes (Cluster B) of ChP in ChP-KO mice reveals RFX consensus motif. G. Enriched GO Biological Process (Cluster A) terms for the upregulated genes in ChP-KO mice. H. The top 10 most differentially upregulated genes at the ChP in ChP-KO mice. I-K. qRT-PCR analysis of inflammation related genes, ICAM1(I), TNFAIP3 (J), GBP4 (K) at the ChP. N = 3 mice/group. *** *P < 0.001*, ** *P < 0.01*, * *P < 0.05*. Data are presented as Mean ± SEM.

Contrastingly, GO analysis and the list of genes that were upregulated in ChP-KO mice showed an increase in immune, defense, and inflammatory responses (Figures 2G and 2H). Among these genes, there was an increased expression of genes associated with innate and adaptive immune responses. To confirm the RNA-seq results, we performed quantitative PCR for the expression of each mRNA. The expression of a wide range of immune- and inflammation-related genes was significantly upregulated in the choroid plexus of ChP-KO mice during embryonic development (Figures 2I–2K and S2F–S2K). These results indicate that CAMDI deficiency in the ChP causes insufficient multiciliogenesis and increased inflammation, suggesting the possibility of functional defects in the choroid plexus.

### Altered Blood-CSF barrier function in CAMDI deficient mice

The blood-cerebrospinal fluid barrier (BCSFB) is an important function of ChP epithelial cells.^42^ To examine whether CAMDI gene deletion causes deficiencies in the barrier function of ChP epithelial cells, we analyzed the expression levels of genes related to cell-cell and cell-ECM adhesion. The mRNA and protein expression levels of ZO-1, a tight junction (TJ)-associated protein, were significantly decreased in CAMDI ChP-KO mice (Figures 3A and 3B). In addition, expressions of TJ-associated Cldn2 and Occludin mRNA were also significantly decreased, suggesting impaired barrier function (Figures S3A and S3B). Consistent with the RNA-seq results showing significantly reduced expression, the expression of Cdhr4, which is associated with intercellular adhesion, was also decreased (Figure S3C). The GO analysis showed that the expression of hemidesmosome-related genes (ITGB4, Col17a1, Plec, and Dsp) were also decreased (Figure S3D; Table S2).

**Figure 3.**
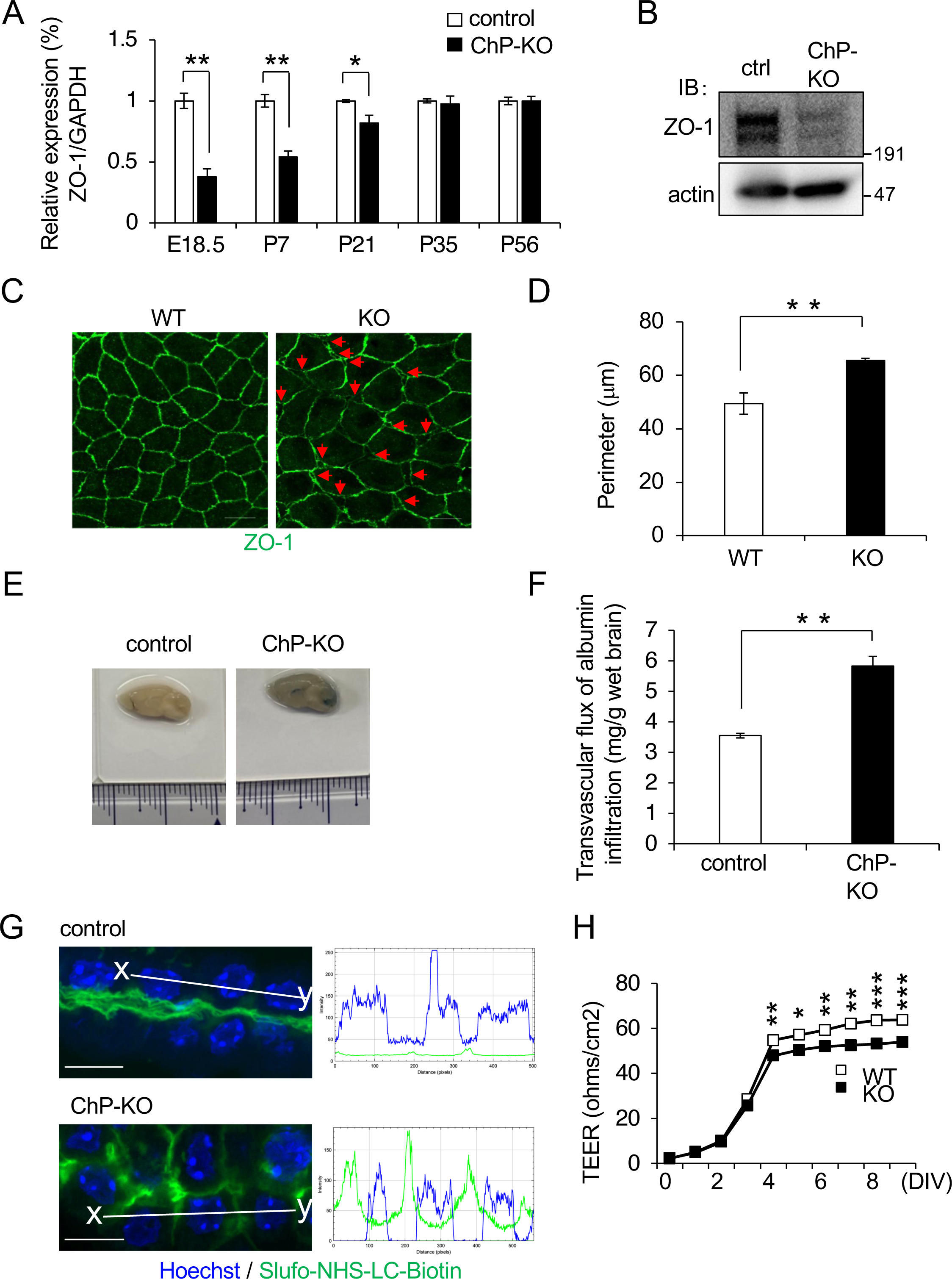
Altered Blood-CSF barrier function in CAMDI deficient mice. A. qRT-PCR analysis of ZO-1 mRNA expression in the ChP of ChP-KO mice (normalized to GAPDH). N = 3 mice/group. B. Western blot analysis of ZO-1 expression in the choroid plexus of ChP-KO mice at E18.5. C. Whole mount immunohistochemical analysis of TJs in CAMDI KO mice at E18.5. ChP were stained for anti ZO-1. Arrows point at membranes that appeared interspaced. Scale bars, 10 μm. D, Cell body perimeter shown in (C). E. Evans blue was injected intraperitoneally into P21 control and ChP-KO mice. After 45 min, the mice were perfused with PBS, and brain permeability was evaluated. F. Evans blue content in the brain was measured at OD620, and the transvascular flux of albumin infiltration is presented after normalization by the wet brain weight. ** *P < 0.01*. G. Slufo-NHC-LC-biotin perfused from the left cardiac ventricle of P21 control and ChP-KO mice. Five minutes after perfusion, frozen sections from the brain were incubated with streptavidin-fluorescein to visualize the distribution of the injected Slufo-NHC-LC-biotin. The ratio line scans along the line (from the x- to y-axis) from the images. Scale bar 10 μm. H. Trans-endothelial electrical resistance (TEER) measured over time in primary CP epithelial cultures from CAMDI-KO mice. N=5 per group. *** *P < 0.001*, ** *P < 0.01*, * *P < 0.05*. Data are presented as Mean ± SEM.

ChP-KO mice received only a single dose of tamoxifen, and therefore contained both gene-knockout and non-knockout cells. To reduce measurement variability, we used whole-body CAMDI KO mice for IHC (Figures 3C and 3D) and BCSFB functional analyses (Figure 3H). IHC analysis of ZO-1 staining revealed the presence of interspaced and large surface areas of ChP epithelial cells in the CAMDI-KO mice (Figures 3C and 3D). These findings suggest that TJs are immature in the ChP due to CAMDI deficiency. Next, we examined the functional role of CAMDI in regulating intercellular adhesion in ChP epithelial cells in vivo. Evans Blue dye was intraperitoneally administered into the blood, and its leakage into the brain parenchyma was quantified.^43^ The brains from ChP-KO mice demonstrated significantly higher leakiness than those from control mice (Figures 3E and 3F). In addition, we tested the in vivo barrier function of TJs by injecting Sulfo-NHS-LC-Biotin, having lower molecular weight than Evans Blue dye, into the perfusion of the heart.^44^ Fluorescence intensity was measured by line-scanning to quantify intercellular leakage, indicating leakage from the blood into the intercellular space between the ChP epithelial cells (Figure 3G). The presence of TJs in the epithelial cells induces trans-epithelial electrical resistance (TEER). To confirm the deficiency in BCSFB function, TEER was measured in primary ChP epithelial cells from CAMDI-KO mice. Compared with WT mice, primary epithelial cells from CAMDI-KO mice showed a significant decrease in TEER (Figure 3H). These results indicate that CAMDI maintains BCSFB integrity and maturation by regulating the expression of TJ and other cell adhesion proteins.

### Delayed social critical period in ChP-KO mice

It has been reported that Otx2 protein from the ChP regulates the beginning and end of the critical period and binocular plasticity.^45^ To examine the effect of CAMDI deficiency in ChP during the critical period, we quantified the expression of Otx2 mRNA and protein. A significant reduction in Otx2 expression was observed in the choroid plexus of ChP-KO mice compared to that in control mice (Figures 4A–4C). An imbalance between the excitatory and inhibitory activities of the nervous system is considered to be a factor in the pathogenesis of ASD.^46^ Otx2 in the ChP has been shown to regulate the expression of proteins in the perineuronal net (PNN) of fast-spiking parvalbumin-expressing GABAergic neurons (PV cells), present in layers III and IV of the cerebral cortex, and is involved in ChP maturation by physically reducing synaptic plasticity by covering PV-positive cells.^47,48^ Therefore, we investigated the numbers of parvalbumin-positive cells (PV+) in the medial prefrontal cortex (mPFC), somatosensory cortex (S1), and primary auditory cortex (A1) at P21. The number of PV+ cells in ChP-KO mice was significantly reduced at mPFC and S1 compared to control mice (Figures 4D and 4E). There was no difference in the number of cells stained for Wisteria floribunba agglutinin (WFA), which recognizes N-acetylgalactosamine, a component of the PNN that surrounds PV+cells (WFA+); however, there was a significant decrease in the number of PV+/WFA+ cells in the mPFC and A1 (Figures 4D, 4E, S4A–S4D).

**Figure 4.**
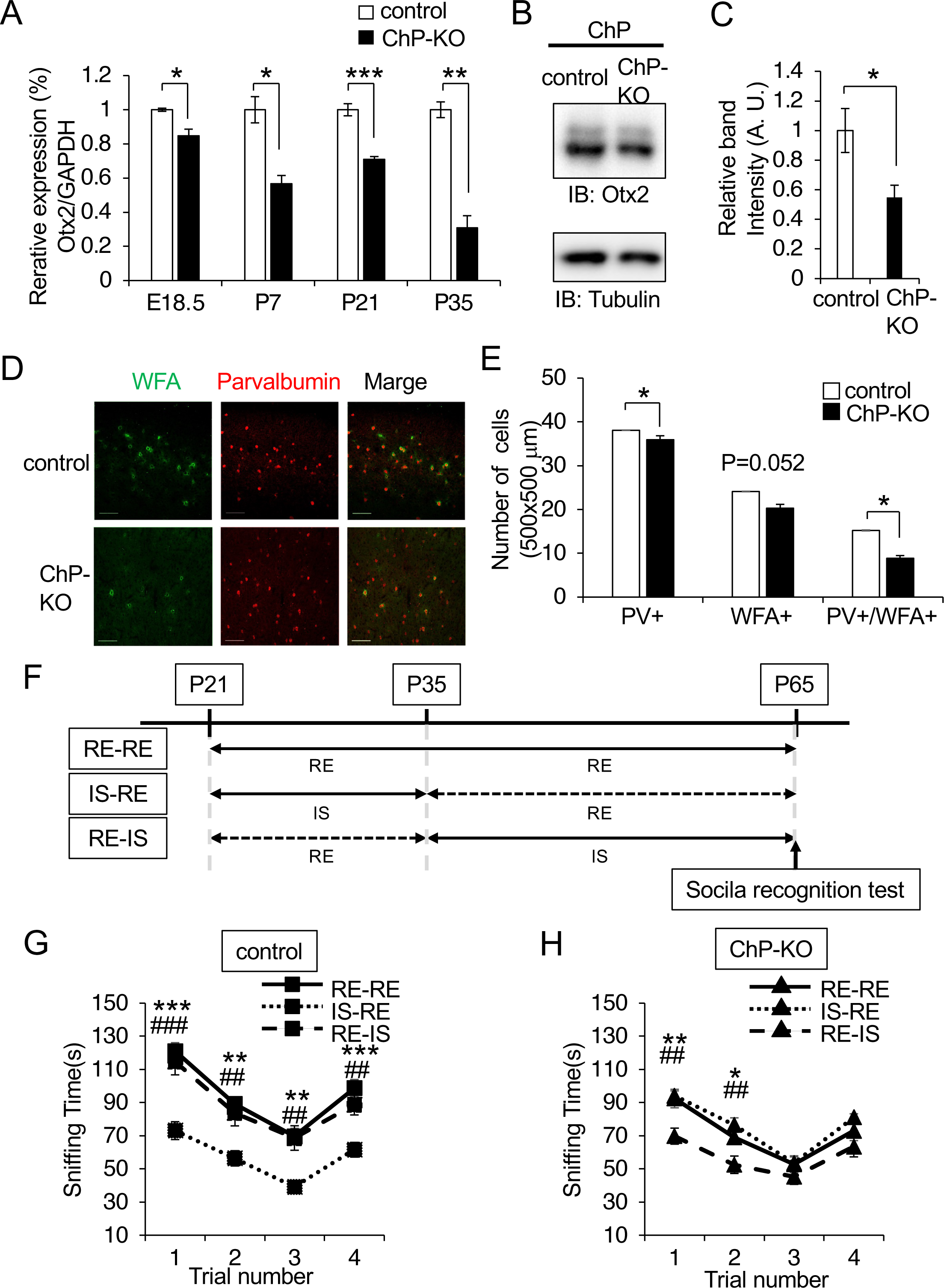
Delayed social critical period in ChP-KO mice. A. qRT-PCR analysis of Otx2 mRNA expression in the ChP (normalized to GAPDH). N = 3 mice/group. B. Western blot analysis of Otx2 expression in the ChP at E18.5. C. Quantification of western blots of Otx2 in the ChP (normalized to Tubulin). N = 3 mice/group. D. Immunohistochemical analysis of PV and WFA in layers II/III of mPFC from P21 control and ChP-KO mice. Scale bar, 100 μm. E. Quantification of PV+, WFA+ and PV+/WFA+ cells in the mPFC. N = 3 mice/group. (500×500 μm area). F. Schema of housing condition for study of critical period. RE-RE: Mice housed in a regular environment (RE, P21 P65) IS-RE: mice housed in an isolated environment (IS, P21–P35) or an RE environment (P36–P65) RE-IS: mice housed in the RE (P21 to P35) and IS (P36 to P65) environments. G. Social recognition times in control mice. Time spent sniffing with stimulus mice. Trials 1–3 represent repeated presentations of the same stimulus mouse, and trial 4 represents the introduction of a novel mouse. N = 16 for RE-RE environment, N = 16 for IS-RE environment, N = 20 for RE-IS environment. * and # indicate significant differences (*P < 0.001* and *P < 0.01*, respectively) between the IS-RE environment and the RE-RE and RE-IS environments. H. Social recognition time in the ChP-KO mice. N = 16, 14, and 15 for the RE-RE, IS-RE, N = 15 for RE-IS environments, respectively. * and # indicate significant differences (*P < 0.01*, and *P < 0.05*) between the RE-IS environment and the RE-RE or IS-RE environments. ***^(###)^ *P < 0.001*, **^(##)^ *P < 0.01*, * *P < 0.05*. Data are presented as Mean ± SEM.

It has been reported that the critical period for social behavior in mice is from P21 to P35.^49^ Thus, we generated three groups of ChP-KO mice under different rearing conditions: group rearing with multiple mice in one cage (regular environment; RE) or isolation (IS) with only one mouse in one cage (RE-RE, IS-RE, RE-IS), and performed a social recognition test on mice reared under these conditions at P65 to evaluate their social cognitive abilities (Figure 4F). As a result, a significant decrease in social recognition, evaluated by sniffing time, was observed in the IS-RE group of control mice, confirming that the social critical period in control mice was between P21 and P35, as previously reported (Figure 4G).^49^ Contrastingly, ChP-KO mice showed significantly reduced social recognition in the RE-IS group, but not in the IS-RE group (Figure 4H). These results indicate that the social critical period is delayed in ChP-KO mice, suggesting that the delay due to immature ChP is a possible factor in ASD-like behavior.

### Metformin improves immature Choroid Plexus and social behavior in ChP-KO mice

Metformin is an oral antidiabetic medication belonging to the class of biguanides. Treatment with metformin is reported to have neuroprotective effects such as reducing inflammation and apoptosis.^50^ Metformin has also been reported to improve blood brain barrier function by restoring the expression of TJ-associated proteins in mouse models for septicemic plague.^51^ To investigate the feasibility of using Metformin for the treatment of ASD-like behaviors, we tested whether oral administration of Metformin improved the social critical period and social behavior in ChP-KO mice during P7 to P21, corresponding to the period just prior to the social critical period (Figure 5A). Despite a slight increase in Otx2 expression, qRT-PCR analysis showed significant recovery of TTR expression in the ChP at P21 in ChP-KO mice treated with Metformin (Figures 5B and S5A). Significant increases in the mRNA levels of TJ-related genes ZO-1 and Cldn2 were observed in Metformin-treated ChP-KO mice (Figures 5C and S5B). In addition, ICAM1 and TNFAIP3 expression significantly decreased in Metformin-treated ChP-KO mice, confirming that inflammation was improved to the same extent as that in control mice (Figures S5C and S5D). To verify whether the barrier function was improved, Evans Blue was administered intraperitoneally in mice at P21. Leakage in Metformin-treated ChP-KO mice improved to the same level as that in control mice (Figure 5D). In vitro transwell assays using primary ChP epithelial cells showed that the leakage to the lower layers observed in ChP-KO cells was significantly improved by Metformin treatment (Figure 5E). These results indicate that Metformin treatment improves the barrier function of ChP epithelial cells, which is vulnerable to CAMDI-deficiency.

**Figure 5.**
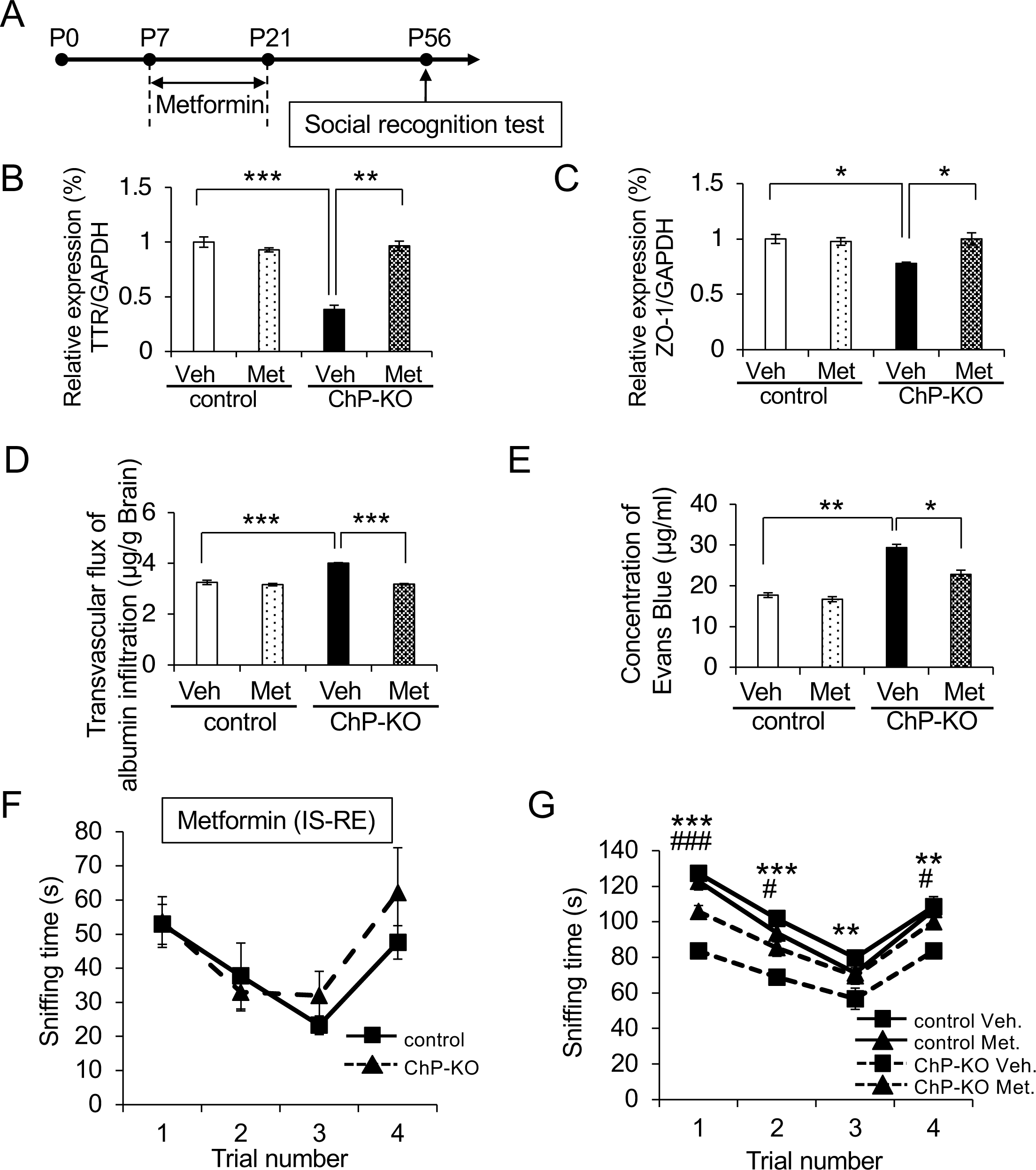
Metformin improves BCSFB functions and social behavior in ChP-KO mice. A. Schematic showing study design: Metformin was administered orally once every day to mice during P7 to P21. B, C. qRT-PCR analysis of TTR (B) and ZO-1 (C) mRNA expression in the ChP at P21 (normalized to GAPDH). N = 3 mice/group. D. Evans blue content in the brain was measured at OD620, and the transvascular flux of albumin infiltration is presented after normalization by the wet brain weight. E. Monolayer permeability was measured using the Transwell-Evans Blue assay. Evans Blue concentration was measured in the lower chambers. F. Social recognition time. Time spent sniffing with stimulus mice. N = 7 for IS-RE environment of Metformin-treated control mice and N = 9 for IS-RE environment of Metformin-treated ChP-KO mice. G. Improvement in social recognition time in Metformin-treated ChP-KO mice. Social recognition time. Time spent sniffing with stimulus mice. N =16 for the RE-RE environment; N = 16 for the IS-RE environment; N = 7 vehicle-treated control mice; N = 9 Vehicle-treated ChP-KO mice; N = 7 metformin-treated control mice; and N = 13 Metformin-treated ChP-KO mice. * indicates significant differences (*P < 0.001* and *P < 0.05*) between the control vehicle treatment and ChP-KO Vehicle treatment. ^#^ indicates significant differences (*P < 0.001*, and *P < 0.05*) between ChP-KO metformin treatment and ChP-KO Vehicle treatment. ***^(###)^ *P < 0.001*, ** *P < 0.01*, *^(#)^ *P < 0.05*; Two-way ANOVA followed by Bonferroni’s post hoc test. Data are presented as Mean ± SEM.

To examine the effects of Metformin administration on the social critical period, we created IS-RE groups and performed a social recognition test at P56. The results showed that Metformin treatment shifted the critical period of ChP-KO mice from P35–56 to P21–35, which coincided with the critical period of control mice (Figure 5F). Furthermore, Metformin-treated ChP-KO mice showed improvement in the social recognition test (Figure 5G). These results indicate that recovering BCSFB function by promoting the maturation of the ChP through Metformin administration before the critical period improves social behavior by normalizing the critical period.

### Immature Choroid Plexus in multicilia-deficient mice and ASD model mice

It has been reported that mice homozygous for targeted insertion of the GFP::CreERT2 cassette into the FoxJ1 locus show a phenotype with multiciliogenesis insufficiency, similar to mice lacking the FoxJ1 gene (Figure 6A).^36^ To confirm that the failure of multiciliogenesis affects ChP maturation, we generated FoxJ1^CreERT2::GFP^ Cre/Cre mice. Although there was no difference in CAMDI mRNA expression (Figure S6A), the expression of the cilia-related gene, Hydin, decreased in Cre/Cre mice (Figure S6B). The expression of ChP differentiation marker genes such as TTR, Otx2, and FolR1, was significantly decreased (Figures 6B, 6C, and S6C). Additionally, the expression of mRNAs encoding TJs, such as ZO-1, Cldn2, and Occludin, significantly decreased (Figures 6D, 6E, and S6D). Furthermore, the mRNA expression of inflammatory factors, such as Tlr2, IL-1γ, ICAM1, IFN-γ, GBP5, and GBP4, was significantly increased in Cre/Cre mice (Figures 6F, 6G, and S6E–S6H), suggesting that inflammation is due to immature ChP with BCSFB dysfunction. These findings indicated that multiciliogenesis is required for ChP maturation.

**Figure 6.**
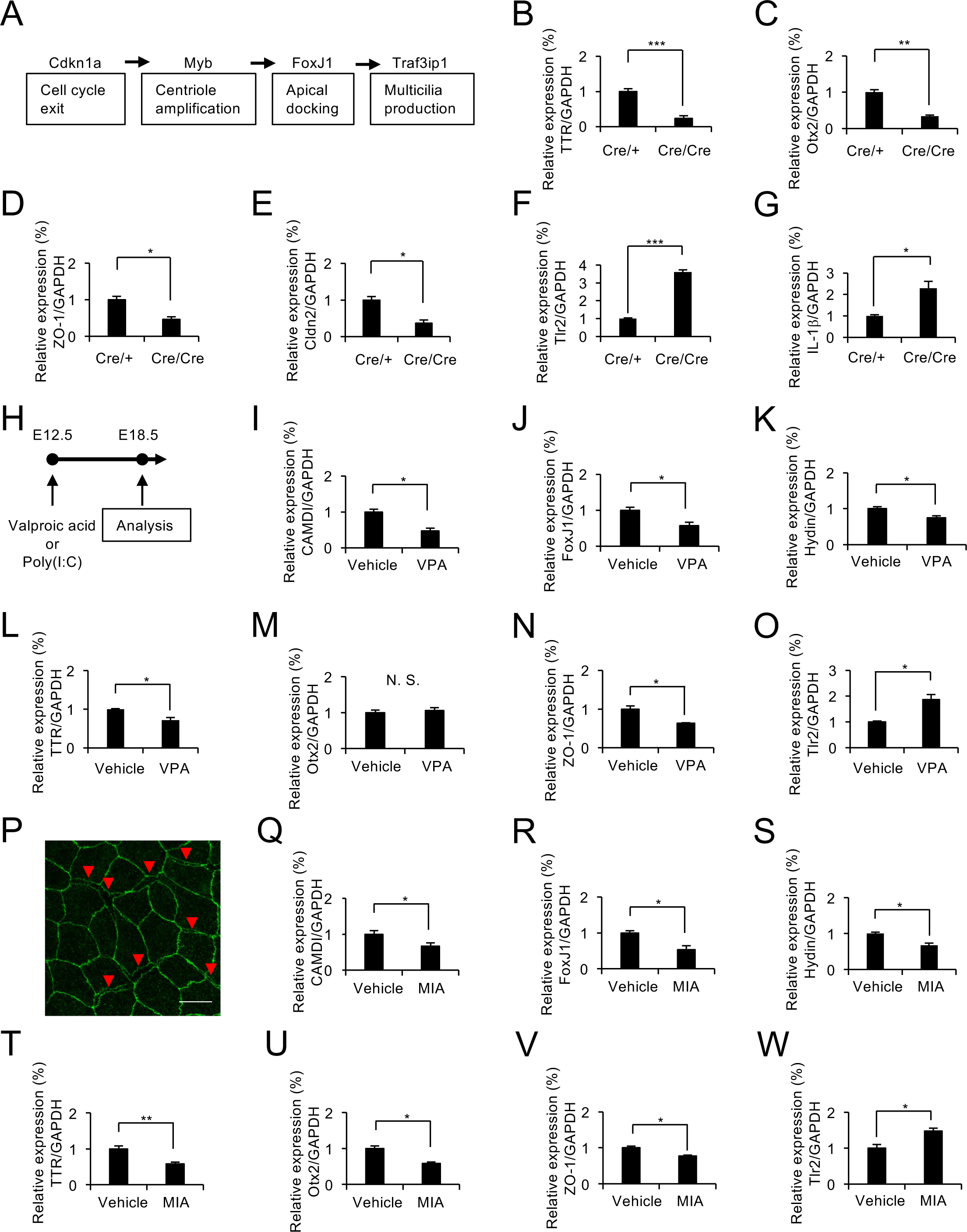
Immature Choroid Plexus in multicilia-deficient and ASD model mice. A. Schematic showing key steps of multiciliated cell differentiation. B–G. qRT-PCR analysis of FoxJ1 Cre/Cre mice, multicilia deficient mice at E18.5. Expression of TTR (B), Otx2 (C), ZO-1 (D), Cldn2 (E), Tlr2 (F), and IL-1β (G). N = 3 mice/group. H. Schematic showing study design: Valproic acid (VPA) or Poly (I:C) for Maternal Immune Activation (MIA) were administered interperitoneally at E12.5. I–O. qRT-PCR analysis of VPA mice at E18.5. Expression of CAMDI (I), FoxJ1 (J), Hydin (K), TTR (L), Otx2 (M), ZO-1 (N), Tlr2 (O). N = 3 mice/group. P. Whole mount immunohistochemical analysis of TJs in MIA mice at E18.5. ChP were stained for anti ZO-1. Arrows point at membranes that appeared interspaced. Scale bars, 10 μm. Q–W. qRT-PCR analysis of MIA mice at E18.5. Expression of CAMDI (Q), FoxJ1 (R), Hydin (S), TTR (T), Otx2 (U), ZO-1 (V), Tlr2 (W). N = 3 mice/group. *** *P < 0.001*, ** *P < 0.01*, * *P < 0.05*. Data are presented as Mean ± SEM.

To examine whether an immature ChP was observed in ASD model mice, we used Valproic acid (VPA) and Maternal Immune Activation (MIA) model mice (Figure 6H).^52,53^ The expression of CAMDI, cilia-related genes (FoxJ1, Hydin), differentiation marker genes (TTR) and TJs genes (ZO-1, Cldn2) was significantly decreased, while that of inflammatory factors (Tlr2, TNFAIP3, IL-1β) was increased in VPA mice (Figures 6I– 6O, S6I-S6L). In addition, we observed that the expression of CAMDI, cilia-related genes (FoxJ1, Hydin), differentiation marker genes (TTR, Otx2) and TJs genes (ZO-1, Cldn2) was significantly decreased, while that of inflammatory factors (Tlr2, TNFAIP3, IL-6, IL-1β) was increased in MIA mice (Figures 6Q–6W, S6M-S6P). IHC analysis of ZO-1 staining revealed the presence of interspaced and large surface areas of ChP epithelial cells in the MIA mice (Figures 6P). These findings indicated that the gene expression patterns of FoxJ1 Cre/Cre mice and ASD model mice were very similar to those of the choroid plexus in ChP-KO mice, suggesting that immature ChP is a common feature of ASD.

### Immature Choroid Plexus in ASD patient-derived organoids

To explore whether an immature ChP with impaired ciliogenesis is a factor in ASD-like behaviors, we used the cilia gene dataset (CiliaCarta^54^ and the ASD-related gene dataset SFARI, https://gene.sfari.org). The Venn diagram showed that 87.2% (975/1118 = 87.2%) of ASD-related genes and 75.7% (708/935 = 75.7) of ciliated genes were expressed in the ChP (Figure 7A; Table S3). Approximately 6.2% (69/1118 = 6.2) of the ASD-associated genes were related to ciliogenesis. Factors common to all three datasets included 65 genes, along with the RFX3 and RFX4 genes. These findings suggest that the onset of ASD may result from immature ChP, including failure of ciliogenesis.

**Figure 7.**
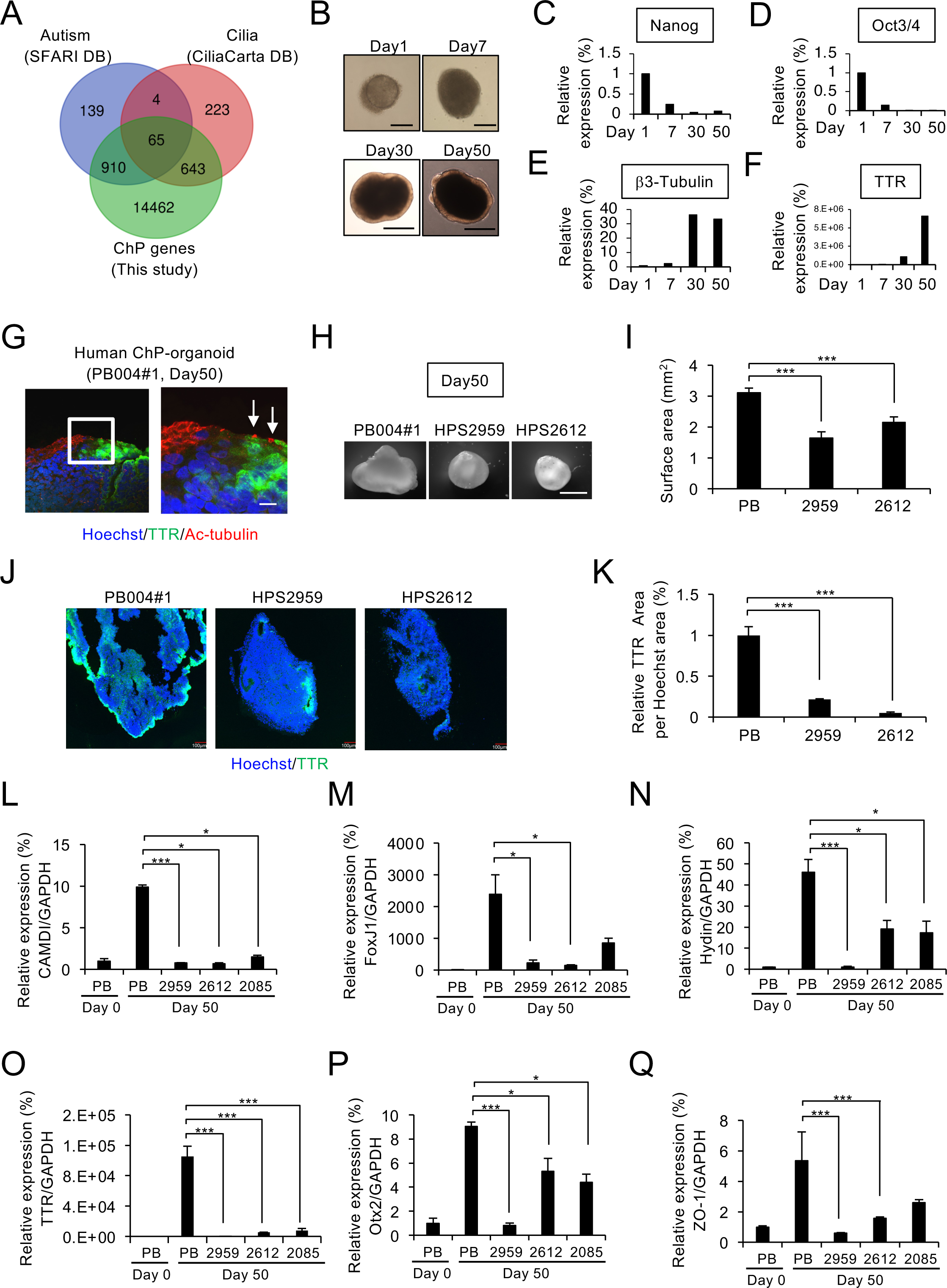
Immature Choroid Plexus in ASD patients-derived organoids. A. Venn diagram showing common and unique sets of Differentially expressed genes between SFARI (ASD, blue), CiliaCarta (Cilia, red) and Choroid plexus RNA sequencing (this study, green) datasets. A total of 65 mRNA are common to all three databases. B. Progression of ChP organoid development from hiPSCs. Sample images at day 1, 7, 10, 30, and 50 in vitro. Scale bars, 200 μm (Day 1, 7), 1000 μm (Day 30, 50). C, D. qRT-PCR analysis of hiPSCs specific marker gene, Nanog (C) and Oct3/4 (D). E. qRT-PCR analysis of ectoderm marker gene, β3-Tubulin. F. qRT-PCR analysis of ChP marker gene TTR. G. Immunohistochemical analysis for TTR (ChP differentiation marker, green), Ac-tubulin (cilia, red) and Hoechst (nuclear, blue) of the ChP organoid at Day 50 (made from control PB004#1 hiPSCs). Scale bar, 10 μm. H. Development of ChP-organoid from ASD patient-derived hiPSCs. Scale bar, 100 μm. I. Quantifications of surface area based on 2D bright field image measurements at Day 50. N = 3 organoids/group. J. Immunohistochemical analysis of TTR (green) from control (PB004#1), ASD patient-(HPS2959, 2612) derived organoids at Day 50. Scale bar, 100 μm. K. Quantitative analysis of TTR area per Hoechst area of Figure 7H. N = 3 organoids/group. L-Q. qRT-PCR analysis of CAMDI (L), FoxJ1 (M), Hydin (N), TTR (O), Otx2 (P) and ZO-1 (Q) mRNA expression in the ChP organoid made from healthy-(PB004#1 refer to as PB) and patient-derived hiPSCs at Day 50, normalized to GAPDH. N = 3 organoids/group. *** *P < 0.001*, ** *P < 0.01*, * *P < 0.05*. Data are presented as Mean ± SEM.

ChP organoids have been previously reported to generate and display a morphology similar to cuboidal epithelial cell-like human ChP epithelium expressing TTR, Otx2, and aquaporin 1 (AQP1). ^55^ Therefore, we generated ChP organoids from healthy human iPS cells (hiPSCs; PB004#1, referred to as PB) (Figure 7B). Quantitative PCR analysis showed a decrease in the expression of the hiPSC marker genes Nanog and Oct3/4 over time (Figure 7C and 7D). In addition, a significant increase in the expression of TTR, a ChP marker gene, was observed despite no difference in the expression of β3-tubulin, a neuroectoderm marker gene, between Day 30 and 50 (Figure 7E and 7F). IHC analysis revealed that cells not expressing TTR showed diffused Ac-tubulin signals, whereas cells expressing TTR showed an aggregated signal on the apical surface, similar to the intrinsic multicilia of ChP epithelial cells (Figure 7G).

To investigate the possibility that the onset of ASD is due to an immature ChP, we generated ChP organoids from ASD (HPS2959, 2612) patient-derived hiPSCs.^56,57^ At the ChP induction stage (Day 30), although TTR expression decreased in HPS2612, no difference in the gene expression levels of the neural progenitor cell marker Pax6, the neuronal marker β3-tubulin, the ChP markers TTR and Otx2 was observed between healthy subjects and ASD patients (Figures S7A–S7F), indicating that the developmental abnormality of the ChP did not occur until the early stages. When organoid size was assessed at the ChP differentiation stage (Day 50), the surface area of patient-derived organoids was significantly reduced by approximately 0.5-fold (Figures 7H and 7I). Quantification of TTR staining areas showed that patient-derived organoids were approximately 20% of the controls and were significantly reduced (Figures 7J and 7K). To further prove the role of immature ChP in ASD symptoms, an analysis with the addition of Angelman syndrome (AS), characterized by developmental delay, lack of speech, motor dysfunction, epilepsy, and autistic features,^58^ was carried out using patient-derived hiPSCs (HPS2085, deletion of 15q11.2-15q11.3). Although there was no significant difference between healthy controls and AS patients specific to FoxJ1 and ZO-1, the expression levels of CAMDI, multiciliogenesis-related genes (FoxJ1 and Hydin, which are involved in cilia motility and one of the 65 genes in Figure 7A; Table S3), ChP differentiation-related genes (TTR, Otx2) and a TJ-related gene (ZO-1) was significantly decreased in ASD and AS patient-derived ChP organoids compared to healthy human iPS cell-derived ChP organoids at Day 50 (Figures 7L–7Q). These results indicate that a ChP with insufficient multiciliogenesis, decreased expression of differentiation genes, and defective TJ formation in BCSFB lead to dysfunction in ASD and AS patient-derived ChP organoids, indicating that immature ChP is a common feature in ASD pathology.

## DISCUSSION

Current research on the biological pathogenesis of ASD is dominated by studies investigating factors in the brain, including neuronal proliferation, differentiation, and migration; spine formation; network formation; E/I balance; inflammation; activation of microglia and astrocytes; and abnormal oligodendrocyte function. However, research has not progressed further despite studies revealing that ChP functionally matures prior to brain development, suggesting its involvement in neuro-developmental disorders, including ASD. We hypothesized that “immature ChP” is a factor in ASD, and were able to show “immature ChP” as a common feature in mouse models of ASD (CAMDI ChP-KO, VPA and MIA) and ChP organoids derived from ASD patients.

### ASD-like behaviors due to immature ChP with multiciliogenesis deficiency

We have previously reported that CAMDI-deficient mice have immature centrioles and exhibit ASD-like behavior.^34^ Comprehensive gene expression analysis revealed decreased expression of genes specific to the ChP, including TTR, and we hypothesized that defective ciliogenesis is associated with immature ChP. Primary cilia signaling has been reported to be involved in the formation of inhibitory neural networks.^59^ Primary cilia are important for the functional maturation of glutamatergic synapses during neurogenesis in adults.^60^ Several studies have indicated the importance of cell lineage determination for differentiation and maturation. Primary cilia formation in the retina is involved in cell maturation, and any abnormalities can disrupt the establishment of the apicobasal polarity leading to development of an immature retinal pigment epithelium.^61,62^ It has been proposed that ciliary dysplasia has a profound effect on brain development.^63,64^ In ChP, multiciliogenesis is required for differentiation and maturation.

It has been reported that ChP tumor cells possess a solitary primary cilium as a result of Notch-mediated suppression of multiciliate differentiation.^39^ These reports suggest that deficiency of ciliogenesis is necessary for the structural and functional maturation of cells and that its disruption may contribute to the onset of several diseases.

FoxJ1 gene is known to be essential for multiciliogenesis.^65^ FoxJ1 Cre/Cre mice, which is equivalent to a deficiency of the FoxJ1 gene, showed decreased expression of differentiation and maturation marker genes such as TTR and Otx2, and TJ-related genes such as ZO-1 and Cldn2, and increased expression of inflammatory marker genes such as IFNγ, IL-1β, and ICAM1. Insufficient formation of BCSFBs, an important function of ChP epithelial cells, is thought to be responsible for inflammation. Both CAMDI ChP-KO and FoxJ1 Cre/Cre mice have multiciliogenesis deficiency and very similar phenotypes, including reduced expression of TTR, Otx2, and ZO-1 alongside increased inflammation. Although multiciliogenesis deficiency alone cannot be definitively stated as a factor in the development of an immature ChP, it is shown to be a contributing factor. Delayed multiciliogenesis may lead to immature ChP during the critical period, resulting in a negative impact on social behavior. In other words, the maturation of the ChP at an appropriate time is necessary for the subsequent development of social behavior. Altogether, these results indicate that multiciliogenesis is essential for the maturation of ChP, suggesting that immature ChP is associated with the onset of ASD-like behaviors.

### Treatment of ASD-like behavior due to maturation of the ChP

The ChP matures prior to brain development. ^66^ The BCSFB is functionally mature from early development prior to the vasculature.^67^ Protein concentrations in the CSF are high immediately before and after birth,^24^ suggesting that maturation of the ChP may be important for the acquisition and establishment of higher cognitive functions. IGF2 is present in the fetal CSF and regulates a proliferative niche for neural progenitor cells,^68^ indicating that a variety of trophic and neurotrophic factors present in the CSF play important roles in brain development from fetal stage to the early postnatal period. One such protein in the CSF is TTR, a differentiation marker of ChP epithelial cells, which binds to thyroid hormones and retinoic acid and is responsible for transporting them from the ChP to the brain via the CSF.^69,70^ Thyroid hormone homeostasis and the expression of downstream genes are altered during pregnancy in ASD patients.^71,72^ The decreased expression of TTR in CAMDI ChP-KO mice suggests that the amount of thyroid hormones and retinoic acid transported into the brain may be reduced, resulting in ASD-like behavior.

Reduced expression of Otx2 was observed in CAMDI ChP-KO mice. Otx2 and TTR are expressed in differentiated ChP epithelial cells with multiple cilia, but not in ChP tumor cells.^39^ Surprisingly, Otx2 is synthesized and transferred from outside the visual cortex to enable visual cortical plasticity.^22^ In addition, ChP Otx2 protein is released into the CSF with its concentration increasing throughout the visual critical period. After reaching the brain parenchyma, Otx2 regulates the visual critical period by contributing to the PNN formation in PV cells.^45^ PNN development and maintenance are necessary for a number of processes within the CNS, including the regulation of GABAergic cell function and termination of the developmental critical period.^10^

In addition to impaired multiciliogenesis and reduced expression of Otx2, we observed a reduction in the number of PV cells with PNN and a critical period failure in social behavior in ChP-KO mice. Moreover, RNA-seq results showed increased inflammation in the ChP of the ChP-KO mice. The ChP is known to function as a BCSFB from early developmenta prior to brain development.^67^ Pups born from Maternal Immune Activation (MIA) mice are known to exhibit ASD-like behaviors via increased expression of IL-17a from the mother and their own IL-17a receptor.^53^ Furthermore, it has been reported that pups derived from MIA mice show increased inflammation in the ChP, including TJ collapse of ChP epithelial cells and activation of macrophages, associated with increased Ccl2 expression.^73^ These immature ChP and ASD-like behaviors observed in CAMDI ChP-KO mice were significantly improved by the administration of Metformin before the social critical period, indicating that treatment to promote ChP maturation early in life might be effective for ASD treatment.

### Maturation of the ChP as a novel therapeutic target for ASD

In addition to the CAMDI ChP-KO mice, MIA and PVA model mice also presented with impaired multiciliogenesis, reduced CSF-secreted proteins (TTR and Otx2), and increased levels of inflammatory mediators. Furthermore, in ChP organoids derived from patients with ASD, we identified the same phenomena, indicating that immature ChP with quantitative changes in these factors may contribute to the development of ASD. Although the scientific validation that abnormalities in the brain are the cause has not been denied, the results of this study advocate for the maturation of the ChP as a novel therapeutic target for ASD.

### Limitations of the study

In the present study, we identified immature ChP in ASD mouse models and patient-derived organoids. It is unclear which molecular pathways are disrupted by multiciliogenesis deficiency that leads to formation of an immature ChP. In addition, many genes associated with multiciliogenesis and ASD, which are expressed in the choroid plexus, are also expressed and function in the brain. Functional disturbances in various brain regions and cell types have been reported in ASD. However, the extent to which the expression of these genes in immature ChP affects brain development is unclear. Further studies are required to understand the mechanisms underlying the development of ASD.

## ACKNOWLEDGMENTS

We thank Asana Yabe, Yuya Sawara, Rika Tadokoro and Tomoyasu Meguro for technical assistance. We thank Dr. Troy Ghashghaei and Dr. Nagendran Muthusamy for Foxj1CreERT2::GFP mice. This study was supported in part by Grants-in-Aid for scientific research from the Ministry of Education, Culture, Sports, Science, and Technology and the Japan Society for the Promotion of Science (to S.Y, T.Y and T.F.). T.F. was supported by Takeda Science foundation. We would like to thank Editage (www.editage.jp) for English language editing.

## AUTHOR CONTRIBUTIONS

Conceptualization: M.T. and T.F.; methodology: M.T. and T.F.; investigation: M.T., Y.S., A.T., K.N., C.S., N.K., A.H., M.O., S.N. and T.F.; formal analysis: M.T. and T.F.; writing – original draft: M.T. and T.F.; writing – review & editing: M.T. and T.F.; visualization: M.T. and T.F.; supervision: S.Y., T.Y. and T.F.; project administration: T.F.; funding acquisition: S.Y, T.Y and T.F.

### Funding and additional information

This work was supported in part by grants-in-aid for scientific research from the Ministry of Education, Culture, Sports, Science, and Technology and the Japan Society for the Promotion of Science (T.F.). T.F. was supported by Takeda Science foundation.

---

**Table S1. Differentially Expressed Genes list of Gene-chip analysis, related to Figure 1**

**Table S2. GO analysis of RNA-seq data, related to Figure 2**

**Table S3. Common expressed genes list between CiliaCarta (Cilia), SFARI (ASD), and ChP RNA-seq (this study) datasets, related to Figure 7**

**Table S4. Sequences of the primers used in this study, related to the STAR Methods**

## STAR METHODS

### Mice

CAMDI flox/flox mice were described previously^34^ and crossed with FoxJ1^CreERT2::GFP^ mice, which was provided from Dr. Troy Ghashghaei.^36^ For cre-mediated excision of ChP-specific CAMDI, single intraperitoneal injection with either cone oil or the Tamoxifen (50 mg per kg bodyweight, Toronto Research Chemicals) at E15.5. In pharmacological experiments, mice received intraperitoneal injections with Metformin (100 mg per kg bodyweight, Tokyo Chemical Industry) or vehicle (saline) between P7 and P21. All animals were maintained under the university guidelines for the care and use of animals. The experiments were performed after securing Tokyo University of Pharmacy and Life Sciences Animal Use Committee Protocol approval.

Postnatal housing conditions were previously described.^49^ To study the effects of rearing conditions, male mice at the time of weaning (postnatal day 21 (P21)) were randomly divided into three different housing conditions: isolation (IS), regular environment (RE). For social isolation, mice were individually housed in standard cages while for regular environment, 4-5 mice were housed in the same standard cage. In addition to regularly housed mice (RE-RE), when necessary, mice were switched between isolation and regular housing as detailed in the text (IS-RE or RE-IS) at P21 to P35 and P35 to P65.

### Valproic acid (VPA) treatment

Pregnant mice were divided into two groups and either received an intraperitoneal injection of valproic acid (SV; 600 mg/kg; Sigma Co) or vehicle (0.9% NaCl, 100 µl/mouse; control group) on E12.5.

### Maternal immune activation (MIA)

On embryonic day 12.5 (E12.5), pregnant dams were single intraperitoneally injected of poly (I:C) (20 mg per kg bodyweight, Potassium salt; Sigma, P9582-5MG) or vehicle solution (NaCl 0.9%).

### Antibodies

Anti-CAMDI antibody was described previously.^25^ Anti-acetyl-tubulin (T6793), anti-a-tubulin (T9026), anti-b-actin (A5441) antibodies were obtained from Sigma-Aldrich. Anti-transthyretin antibody (NBP1-50256) was obtained from Novus. Anti-Otx2 antibody (ab183951) was obtained from Abcam. Anti-ZO-1 antibody (61-7300) was obtained from Thermo Fisher scientific. Anti-PV antibody (MAB1572) was obtained from Merck Millipore. Wisteria Floribunda Lectin, Fluorescein was purchased from Vector laboratories. Secondary antibodies conjugated with Alexa Fluor 488, and 594 were obtained from Invitrogen. Hoechst 33258 was obtained from Nacalai Tesque.

### RNA isolation from mouse ChP tissue

Total RNA of lateral ventricle choroid plexus tissue was extracted using RNeasy (Qiagen) in accordance with the manufacturer’s instructions. Integrity of the RNA samples was determined by electropherogram analysis in a 2100 Bioanalyzer (Agilent Technologies).

### DNA-chip analysis

Briefly, 1 mg total RNA was analyzed for mRNA profiling using microarray, 3D-Gene® Mouse Oligo chip 24K (Toray Industries Inc., Tokyo, Japan) according to the manufacturer’s protocol. Hybridization signals derived from Cy3 and Cy5 were scaned and the expression level of each oligo was globally normalized using the background-subtracted signal intensity of the entire mRNAs in each array.

### RNA-seq analysis

RNA-seq was performed using choroid plexus from control and ChP-KO mice. For quality control, in order to remove technical sequences, including adapters, polymerase chain reaction (PCR) primers, or fragments thereof, and quality of bases lower than 20, pass filter data of fastq format were processed by Cutadapt (V1.9.1) to be high quality clean data. For mapping, firstly, reference genome sequences and gene model annotation files of relative species were downloaded from genome website, such as UCSC, NCBI, ENSEMBL. Secondly, Hisat2 (v2.0.1) was used to index reference genome sequence. Finally, clean data were aligned to reference genome via software Hisat2 (v2.0.1). For expression analysis, in the beginning transcripts in fasta format are converted from known gff annotation file and indexed properly. Then, with the file as a reference gene file, HTSeq (v0.6.1) estimated gene and isoform expression levels from the pair-end clean data. Differential expression analysis used the DESeq2 Bioconductor package, a model based on the negative binomial distribution. the estimates of dispersion and logarithmic fold changes incorporate data-driven prior distributions, Padj of genes were setted <0.05 to detect differential expressed ones. For GO enrichment analysis, GOSeq(v1.34.1) and iDEP.96 were was used identifying Gene Ontology (GO) terms that annotate a list of enriched genes with a significant padj less than 0.05.

### Quantitative reverse transcription PCR (qRT-PCR)

The reverse transcription reaction was performed using the RevarTra Ace qPCR RT kit (TOYOBO). A quantitative RT-PCR was carried out with a Rotor-Gene Q (QIAGEN) using THUNDERBIRD Next SYBR qPCR mix (TOYOBO) according to the manufacturers’ instructions. qRT-PCR was performed in technical duplicate for per target gene and sample were averaged. At the end of the assay, a melting curve was constructed to evaluate the specificity of the reaction. Gene expression is shown as fold change relative to control mice, following normalization to housekeeping gene GAPDH expression. All primers used in this study are listed in Table S4.

### Western blotting

ChP tissue was lysed in Nonidet P-40 lysis buffer (20 mM Tris HCl, pH 7.2, 2 mM EDTA, 0.5% Nonidet P-40, 8% sucrose, 80 mM dithiothreitol). The lysate was clarified by centrifugation at 15,000g for 10 min and subjected to SDS-PAGE and transferred to polyvinylidene difluoride membranes (Immobilon P; Millipore). Membranes were blocked for 1 h at room temperature (RT) in 5% skim milk in PBST with gentle shaking and incubated with primary anti-bodies overnight. After washing the membranes three times with PBST, they were incubated with the secondary antibody conjugated to horseradish peroxidase for 1h at RT. The blotted membranes were developed using the Immobilon Western chemiluminescent HRP substrate (Millipore) according to the manufacturer’s instructions.

### Immunohistochemistry

For immunohistochemical analysis, brains were fixed by immersion in 4% paraformaldehyde in 0.1 M PBST (PBS containing 0.1% Tween-20), cryoprotected in 20% sucrose, frozen in OCT compound. Coronal sections (20 μm) were cut with a cryostat and were blocked for 1h at RT in PBS containing 5% horse serum and then incubated overnight with the primary antibody, followed by incubation with secondary antibody. The staining was analyzed by confocal microscope (Olympus FV1000-D).

### Evans blue (EB) assay

EB assay was described previously.^43^ Briefly Control and CAMDI ChP-KO mice were anesthetized and Evans blue (40 mg/kg) were injected intraperitoneally (i.p.) for 45 min. Mouse brains were excised, weighed, homogenized in 500 ml PBS, and extracted in 1 ml formamide for 24 h at 60 °C. Evans blue content in the formamide extract was determined by OD_620_ and OD_740_. We used the following equation: corrected 620 nm optical density = (OD_620_ (Evans blue) − (1.426 × OD_740_ (hemoglobin) + 0.03).

### Tracer experiments

Tracer experiments was described previously.^74^ The following solutions were perfused from the left cardiac ventricle of P21 mice. 15 ml of 0.5 mg/ml EZ-LinkTM Sulfo-NHS-LC-Biotin (557 D; Pierce) in PBS. 5 min after perfusion, the whole brain was removed, fixed with 4% PFA, and frozen using liquid nitrogen. In the frozen sections, the distribution of EZ-LinkTM Sulfo-NHS-Biotin were detected by incubation with secondary antibody and/or FITC-Streptavidin.

### Primary culture of ChP cells

Cell cultures were described previously.^75^ Briefly, ChP tissues were dissected and digested with 0.25% trypsin, and manually dissociated into a single-cell suspension (1-2×10^5^ cells/mL). After 2 h non-attached cells were taken and seeded to laminin-coated dish, cover glass or transwell filters (for TEER measurement, and permeability assay). Cells were cultured in Dulbecco’s modified Eagle’s medium (DMEM) supplemented with 10% fetal calf serum (FCS), penicillin-streptomycin.

### TEER measurement

TEER measurement was described previously.^75^ Briefly, ChP cells were plated on 0.4μm pore size, 24well Thincert-TC incert (Greiner BIO-ONE) and analyzed in triplicate by Millicell ERS-2 system (Millipore) in accordance with the manufacturer’s instructions.

### Transwell Evans Blue monolayer permeability assay

Transwell Evans Blue monolayer permeability assay was described previously.^76^ Transwell inserts (Thincert-TC incert, 24well pore size 3.0μm, Greiner BIO-ONE) were used for the permeability assay. Primary ChP cells were seeded on Transwell inserts at a concentration of 1 × 10^5^ cells/well and incubated for 72 h in a confluent cell monolayer. Cells were incubated serum-free medium with vehicle or 2mM Metformin for 24 h. Evans blue-conjugated albumin (final concentration: 0.67 mg/ml) was prepared by diluting a stock solution of 2% Evans blue in a 4% BSA to eliminate any free EB. 100 μl of EB-conjugated albumin was then added to the upper chamber, and 500 μl of 4% BSA was added to the lower chamber. After incubating for 1 h, the liquid in the lower chamber was collected. Absorbance was determined at 620 nm using a micro-plate reader.

### Behavioral study

Male mice were used for behavior studies. Behavior tests were performed as previously report.^34^ The experimenters were blind of the genotype of the tested animals for data collection and analyses.

### Social recognition test

The testing apparatus was a white, plastic transport box (40×40×15 cm). Test mouse was placed in the box and allowed habituate for 10 min, after which a visitor mouse was introduced into the box. After 5 min, visitor mouse was removed. After a 10 min inter-exposure interval, the same stimulus mouse was reintroduced. We repeated this sequence for three trials. In a fourth ‘dishabituation’ trial, the test mice were then exposed to a novel stimulus mouse for 5 min. The time the test mouse spent interacting with each visitor mouse was recorded. Social recognition was measured by social interaction time.

### hiPSCs culture

PB004#1 control hiPSCs^77^ and disease-specific iPS cells^56^ were established from the peripheral blood mononuclear cells using human *OCT4*, *SOX2*, *KLF4* and *MYC* vectors. We used iPS cell lines derived from ASD (HPS2959, HSP2612) and Angelman syndrome (HSP2085) patients provided by the RIKEN BRC through the National BioResource Project of the MEXT, Japan.

### ChP Organoid Culture

ChP Organoid Culture was previously described.^55^ Briefly, undifferentiated hiPSCs grown in feeder-free conditions were detached and replaced with StemFit medium supplemented with 10 μM Y-27632. Embryoid body (EB) were transferred to neural induction medium (DMEM:F12 supplemented with 20% KnockOut Serum Replacement, 1X MEM-NEAAs, 1X GlutaMax, 1X Penicillin-Streptomycin, 1X 2-mercaptoethanol, 0.5 μM LDN-193189, 5 μM SB-431542, 1 μM IWP-2, and 10 μM Y-27632). On day 8, EBs were transferred to ChP induction medium (DMEM:F12 supplemented with 1X N2 supplement, 1X Penicillin/Streptomycin, 1X Non-essential Amino Acids, 1X GlutaMax, 3 μM CHIR-99021, and 200 ng/mL human recombinant BMP-7) and cultured on an orbital shaker rotating at 110 rpm. On day 30, EB were transferred to differentiation medium (1:1 mixture of DMEM/F12 and Neurobasal supplemented with 1X N2 supplement, 1X B27 supplement w/o vitamin A, 1X Penicillin/Streptomycin, 1X Non-essential Amino Acids, 1X GlutaMax, 10 ng/mL BDNF, and 10 ng/mL GDNF). Differentiation medium was replaced every three days.

### Statistical analyses

Statistical significance was determined using unpaired two-tailed Student’s t-tests to compare two groups. For multiple comparison analysis, one-way ANOVA followed Tukey-Kramer post hoc test or Two-way ANOVA followed by Bonferroni’s post hoc test was used. All results are expressed as the mean ± SEM.

### Data and Software Availability

Microarray data is deposited as a MIAME compliant study in NCBI’s Gene Expression Omnibus and are accessible through GEO Series accession number GSE60278 (https://www.ncbi.nlm.nih.gov/geo/query/acc.cgi?token=wlunscusfruhhyr&acc=GSE60278). The raw data of RNA-seq analysis have been deposited in the DNA Data Bank of Japan (DDBJ) Sequence Read Archive under accession number DRA016191.

## Supplemental Information

**Figure S1.**
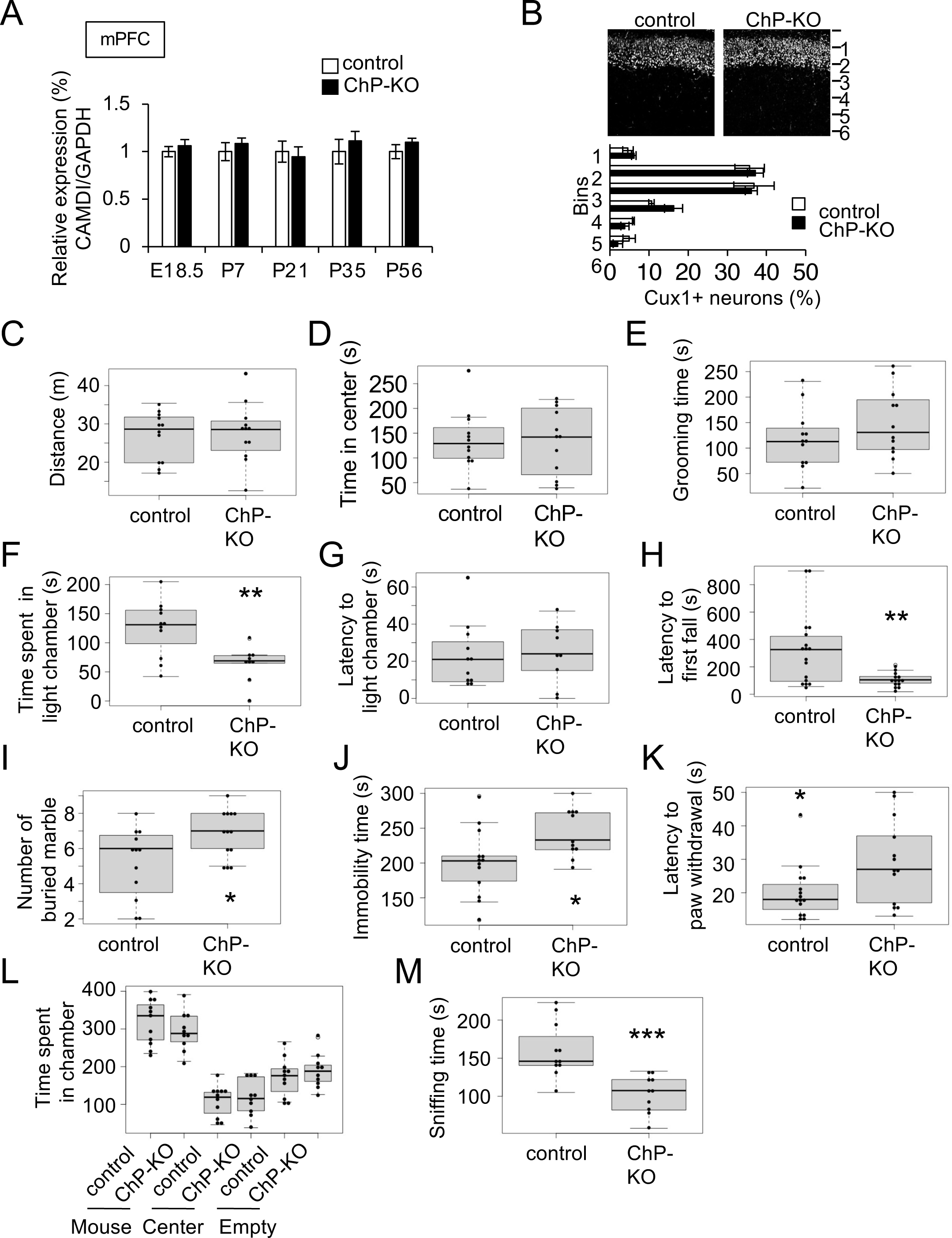
ASD-like behavior in ChP-KO mice, related Figure 1. A. qRT-PCR analysis of mPFC from ChP-KO mice. Expression of CAMDI at ChP (upper panel) and mPFC (lower panel). N = 3 mice/group. B. Immunohistochemical analysis of Cux1-positive neurons in somatosensory cortex at P2 (upper panel). Quantification of the number of Cux1-positive neurons (lower panel). *n* = 3 mice/genotype (control = 913 cells, ChP-KO = 1,531 cells). C–E. Open-field test. Distance traveled (C), time in center (D) and grooming time (E). F, G. Light–dark test. Time spent in light chamber (F) and latency to light chamber (G). H. Cliff avoidance test. I. Marble-burying test. J. Forced swimming test. K. Hot plate test. L, M. Three-chamber social interaction test. The time spent in chamber (L) and the time spent sniffing with stranger mouse (M). Control mice (N = 11) and ChP-KO mice (N= 12). *\*\*\* P < 0.001, ** P < 0.01, * P < 0.05*. Data are presented as Mean ± SEM.

**Figure S2.**
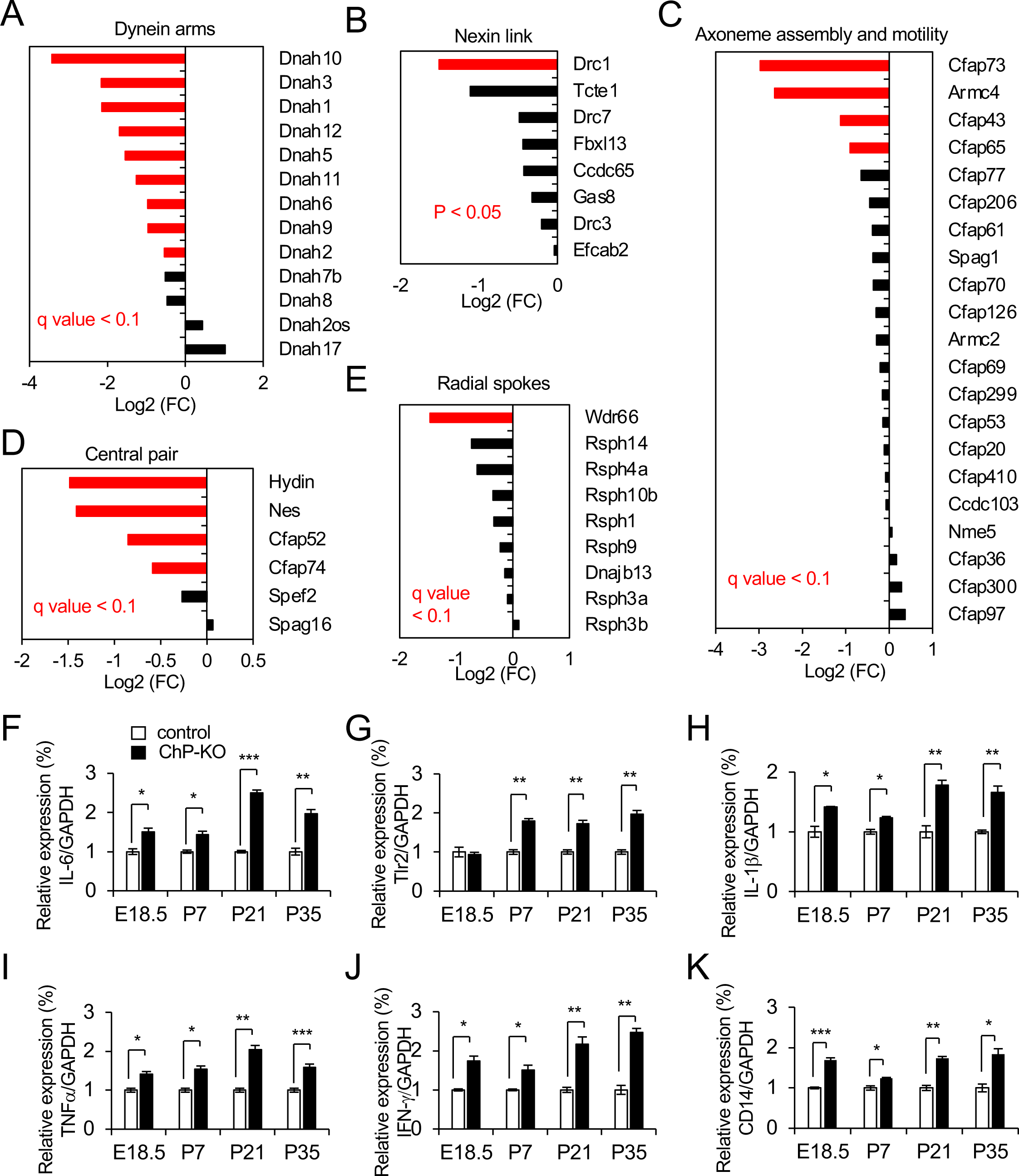
Multiciliogenesis deficiency and increased inflammation in ChP-KO mice, related Figure 2. A–E. The expression of the structural components specific to motile cilia in ChP-K mice. Dynein arms (A), Nexin link (B), Axoneme assembly and motility (C), Central pair (D), Radial spokes(E). F–K. qRT-PCR analysis of inflammation related genes, IL-6 (F), Tlr2 (G), IL-1β (H), TNF-α (I), IFN-γ (J) and CD14 (K) at the ChP. N = 3 mice/group. *\*\*\* P < 0.001, ** P < 0.01, * P < 0.05*. Data are presented as Mean ± SEM.

**Figure S3.**
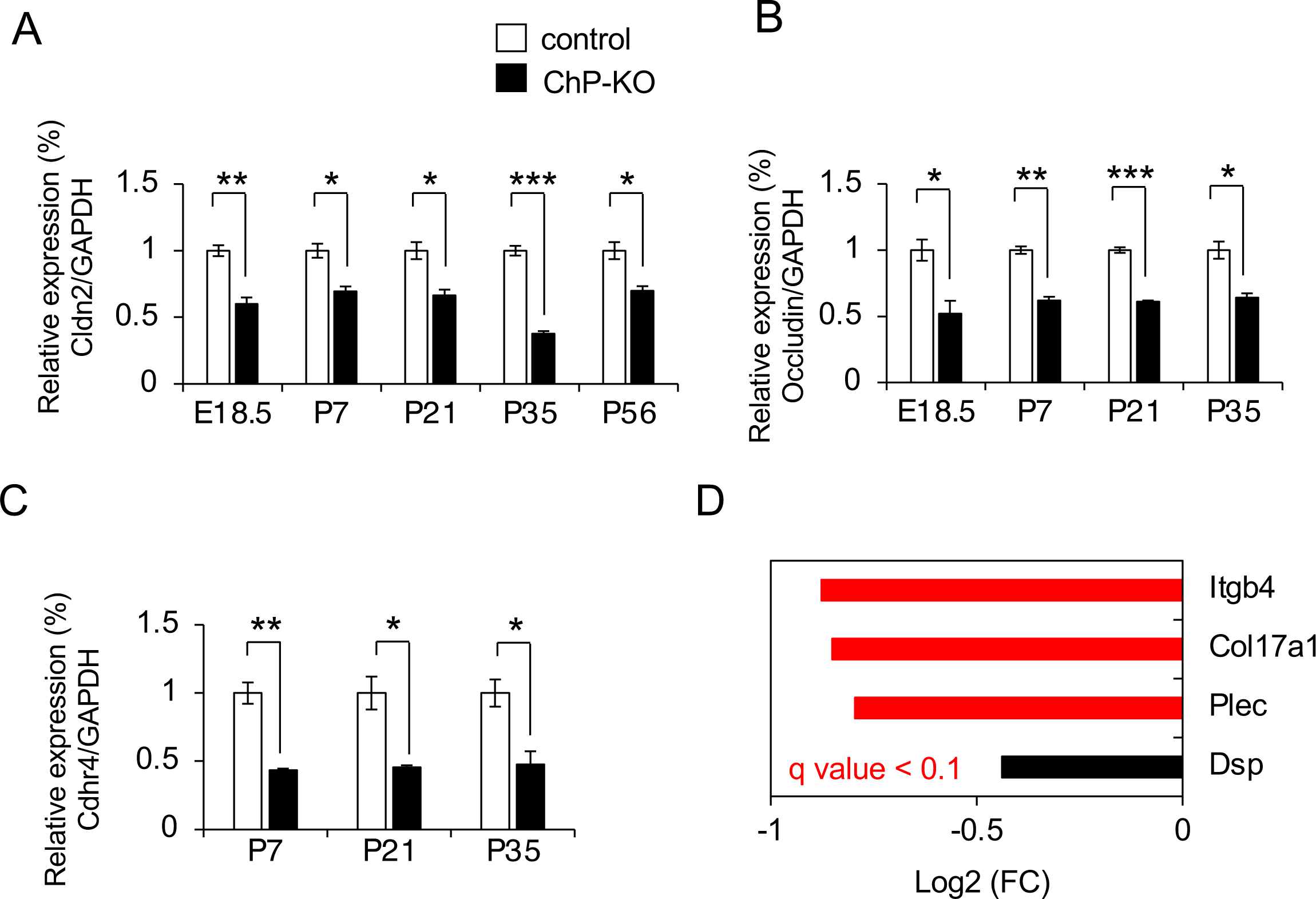
Reduced expression of cell adhesion genes in ChP-KO mice, related Figure 3. A–C. qRT-PCR analysis of inflammation factors in ChP-KO mice compared to control mice. Expression of Cldn2 (A), Occludin (B) and Cadherin Related Family Member 4, Cdhr4 (C). N = 3 mice/group. D. The expression of the Hemidesmosome-related genes in ChP of ChP-KO mice at P21. *\*\*\* P < 0.001, ** P < 0.01, * P < 0.05*. Data are presented as Mean ± SEM.

**Figure S4.**
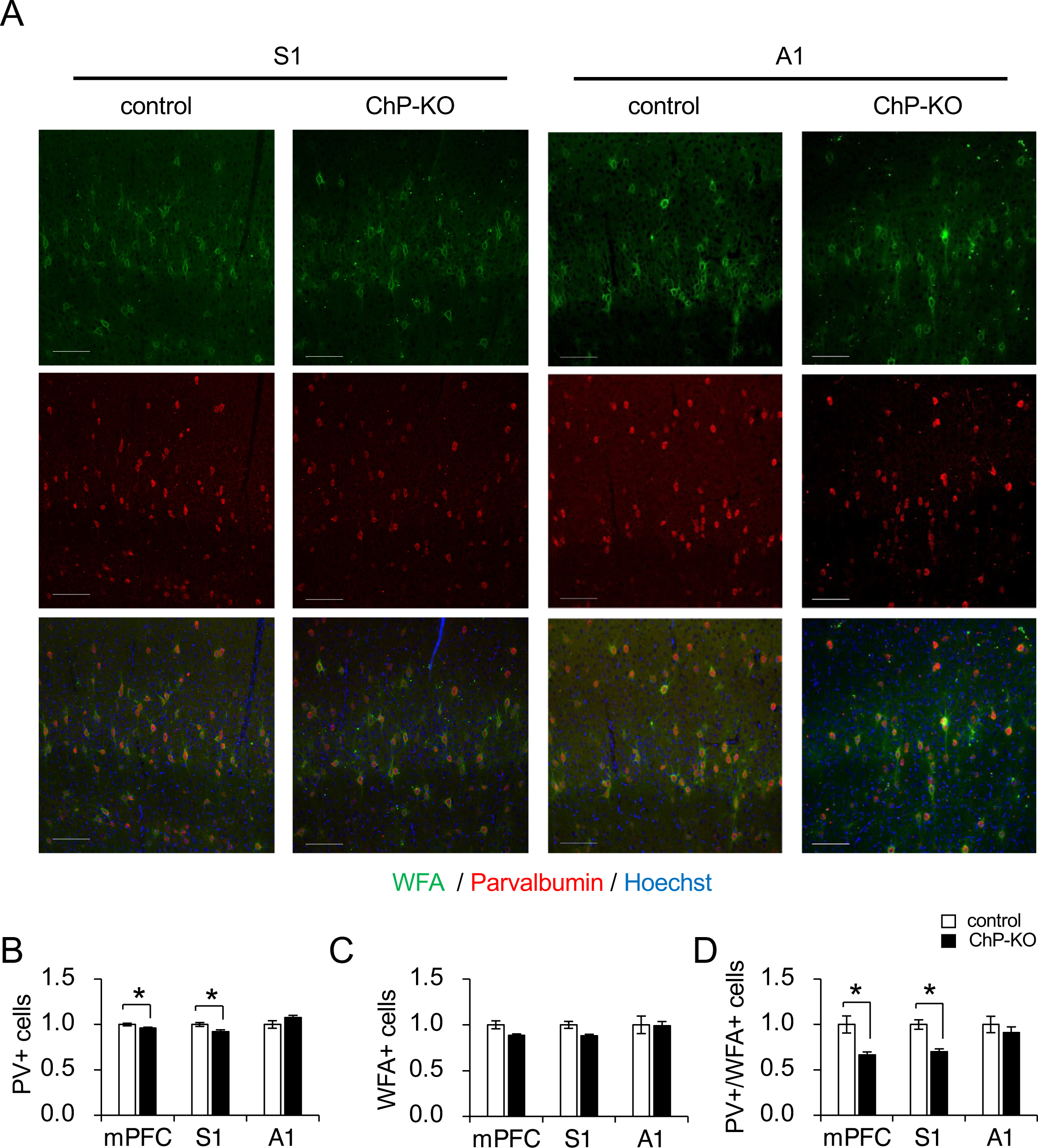
Decrease of PV/WFA cells in ChP-KO mice, related Figure 4. A. Immunohistochemical analysis of Parvalbumin (PV) and WFA in somatosensory (S1), and auditory (A1) from P21 control and ChP-KO mice. Scale bar, 100 μm. B–D. Quantification of PV (B), WFA (C) and PV/WFA (D) positive cells. N = 3 mice/group. *\* P < 0.05.* Data are presented as Mean ± SEM.

**Figure S5.**
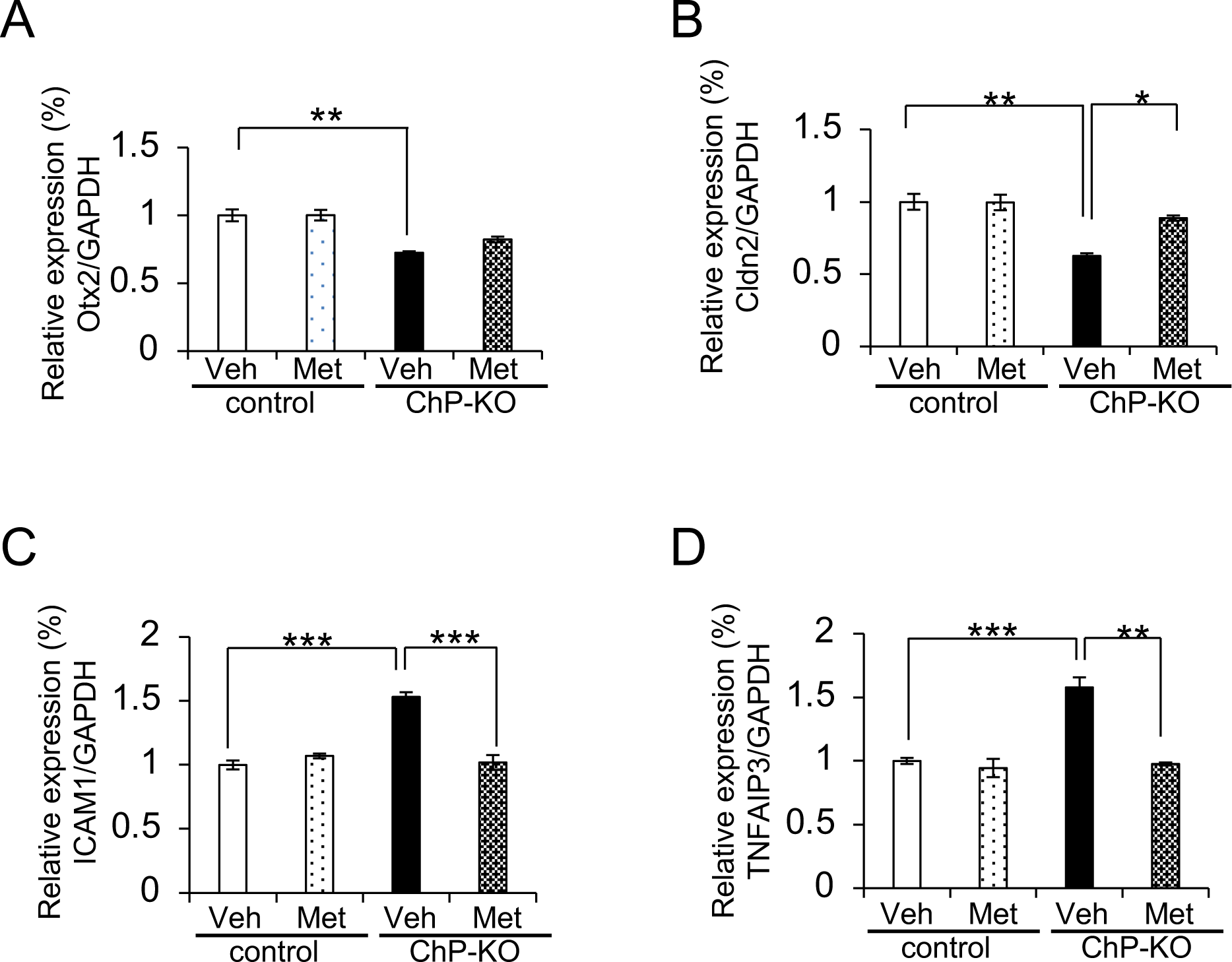
Metformin improves immaturation and inflammation in ChP-KO mice, related Figure 5. A–D. qRT-PCR analysis of Otx2 (A), Cldn2 (B), ICAM1 (C), and TNFAIP3 (D) mRNA expression in the ChP (normalized to GAPDH). N = 3 mice/group. *\*\*\* P < 0.001, ** P < 0.01, * P < 0.05*; Two-way ANOVA followed by Bonferroni’s post hoc test. Data are presented as Mean ± SEM.

**Figure S6.**
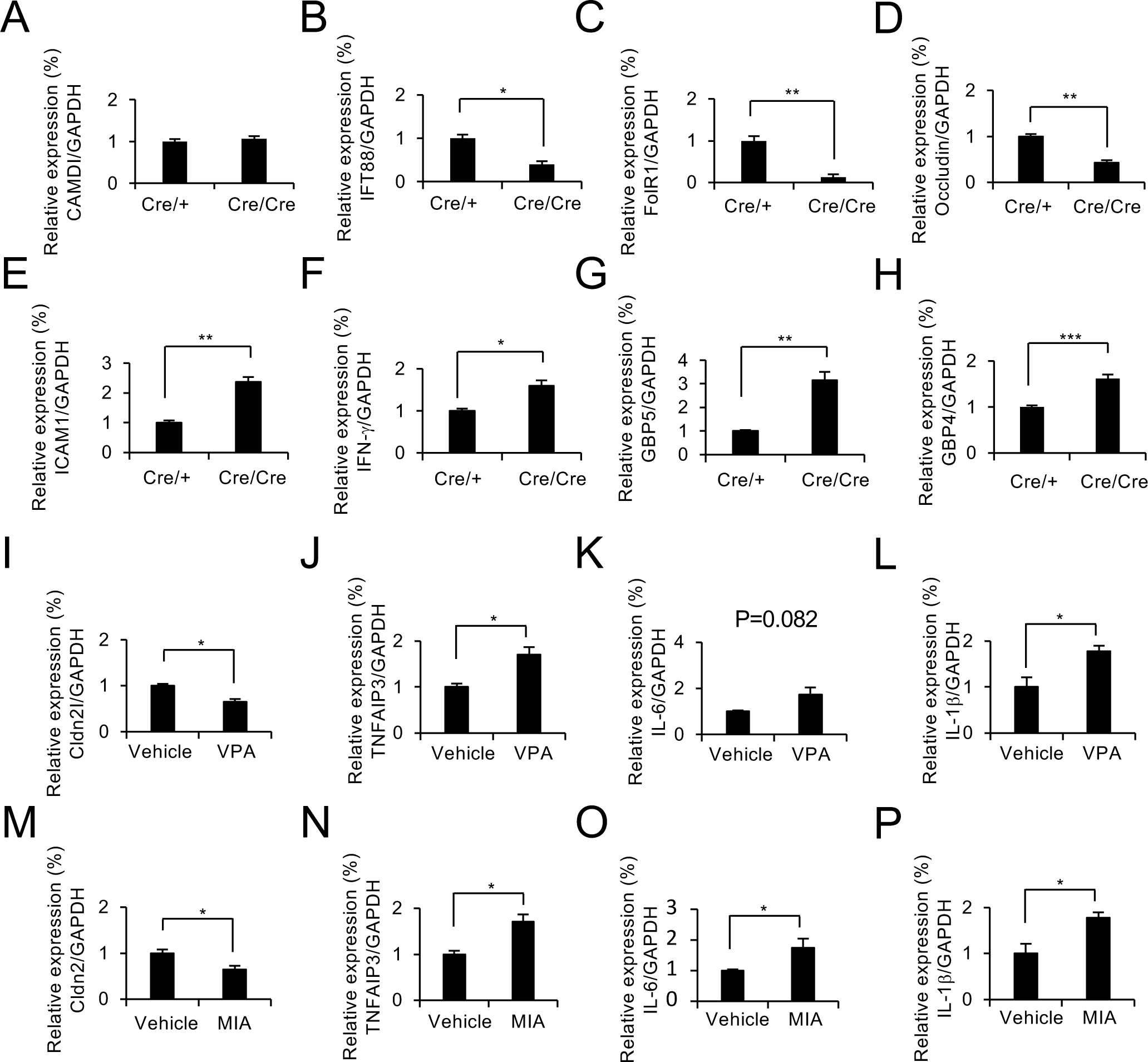
Immature ChP in FoxJ1 Cre/Cre and ASD mice, related Figure 6. A–H. qRT-PCR analysis of FoxJ1 Cre/Cre mice at E18.5. Expression of CAMDI (A), IFT88 (B), FolR (C), Occludin (D), ICAM1 (E), IFN-γ (F), GBP5(G) and GBP4 (H). N = 3 mice/group. I–L. qRT-PCR analysis of VPA mice at E18.5. Expression of Cldn2 (I), TNFAIP3 (J), IL-6 (K), IL-1β (L). N = 3 mice/group. M–P. qRT-PCR analysis of MIA mice at E18.5. Expression of Cldn2 (M), TNFAIP3 (N), IL-6 (O), IL-1β (P). N = 3 mice/group. *\*\* P < 0.01, * P < 0.05*. Data are presented as Mean ± SEM.

**Figure S7.**
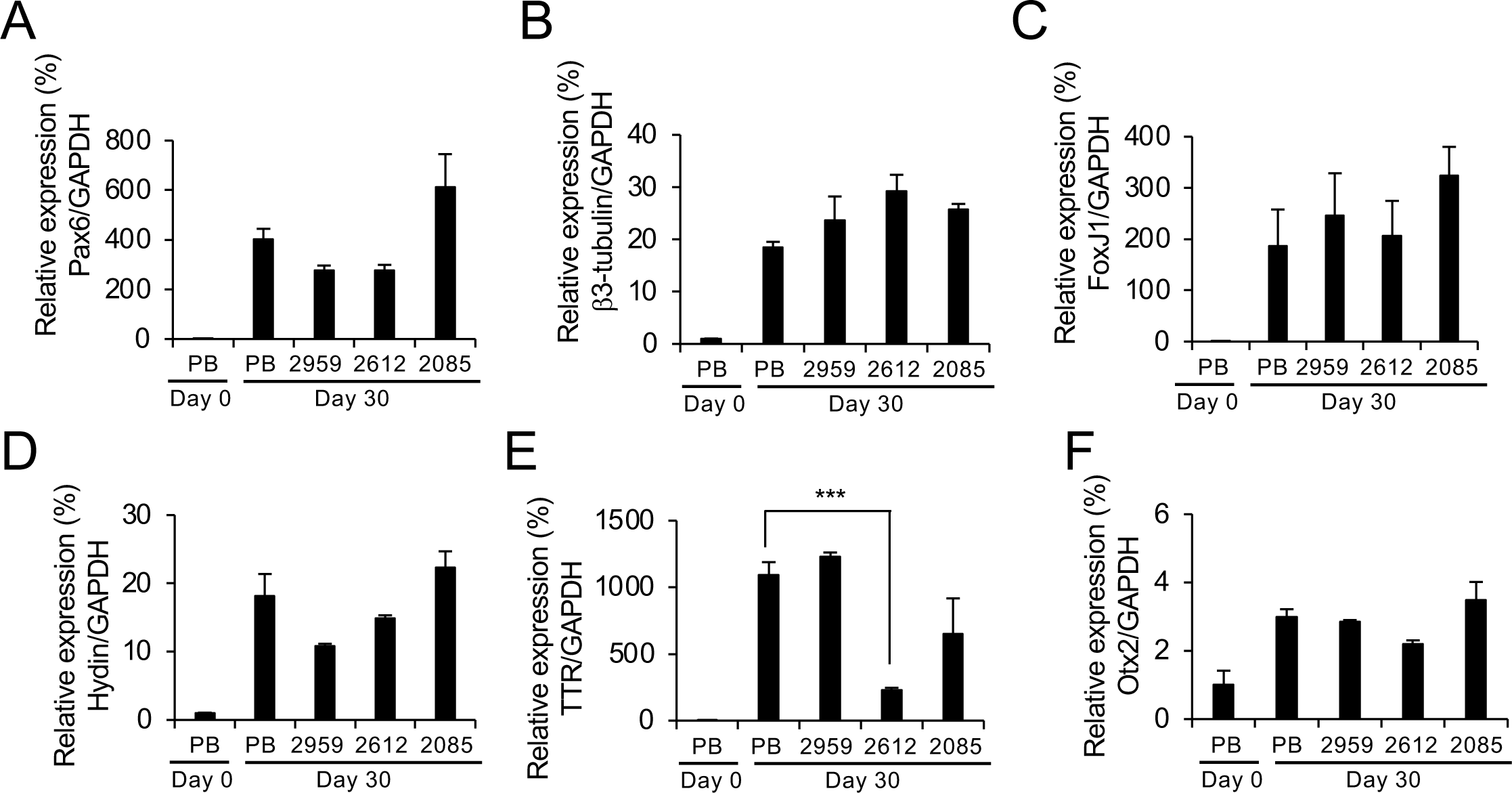
Immature ChP organoids from ASD and AS patients-derived hiPSCs, related Figure 7. A–F. qRT-PCR analysis of Pax6 (A), β3-tubulin (B), FoxJ1 (C), Hydin (D), TTR (E) and Otx2 (F) mRNA expression ChP organoids from ASD patients at Day 30 (normalized to GAPDH). N = 3 organoids/group. *\*\*\* P < 0.001, ** P < 0.01, * P < 0.05*. Data are presented as Mean ± SEM.

